# The Genomic Legacy of the Norman Conquest in Rural England

**DOI:** 10.64898/2026.04.10.716983

**Authors:** Flavio De Angelis, Elizabeth A. Nelson, Sam Leggett, Kalina Kassadjikova, Tanya R. Pelayo, Rob Poulton, Todd C. Rae, Lars Fehren-Schmitz, Lia Betti, Carlos Eduardo G. Amorim

## Abstract

The Norman Conquest of 1066 CE reshaped the political and cultural landscape of England, yet its demographic consequences remain poorly understood, particularly outside elite and urban contexts where historical evidence is concentrated. Here, we investigate the population history of a rural English community spanning the Conquest using genome-wide ancient DNA from the Priory Orchard site, a cemetery in Godalming (Surrey) in use between the 9^th^ and early 13^th^ centuries CE. We generated genomic data from 78 individuals and established radiocarbon dates for 98 individuals from the site. Population genetic analyses place Priory Orchard individuals within the genetic continuum of early medieval populations from the North Sea region. Ancestry modelling indicates that this rural community carried substantial Scandinavian/Viking-related ancestry alongside a persistent Saxon-related component and a smaller French-related contribution. However, stratifying individuals by date, before and after 1066 CE, reveals no clear genome-wide discontinuity across the Conquest horizon, suggesting demographic continuity through this crucial political and social transition. This pattern is consistent with historical and archaeological evidence indicating that many of the most visible transformations following the Conquest occurred primarily among the elite. Our results provide the first genomic perspective on communities living through the Norman Conquest and indicate that rural southern England saw persistent migration links with other areas facing the North Sea rather than abrupt population replacement.

## INTRODUCTION

The centuries preceding the Norman Conquest of 1066 CE represent a formative period in the social, political, and evolutionary history of England. Between the 5^th^ and 11^th^ centuries, repeated episodes of migration and interaction across the North Sea and English Channel reshaped the demographic, cultural, and genetic landscape of the region.^1–6^ Genome-wide studies have confirmed the substantial contribution of the *Adventus Saxonum* to the English gene pool,^1^ while also revealing a complex and regionally heterogeneous process of population movement and integration, reflected in local genetic, isotopic, and archaeological variability within early medieval cemeteries.^6,7^

Archaeological evidence underscores sustained connections among the Atlantic Archipelago, Scandinavia, and Frankish territories throughout the first millennium CE, highlighting the North Sea as a major conduit of mobility and exchange.^8–11^ Isotopic data further demonstrate continued population movement into and within Britain in the centuries leading up to the Norman invasion.^6,12^ These dynamics intensified during the Viking Age, beginning in the mid-8^th^ century, when Scandinavian groups established settlements across Britain and Ireland. Following the arrival of the so-called Great Army in 865 CE, eastern and northern England were incorporated into the Danelaw, with tens of thousands of settlers estimated to have arrived by the end of the 10^th^ century.^13,14^

Recent genomic surveys of Viking-Age populations have shown that “Viking” identity was primarily cultural rather than indicative of a genetically-defined population, with individuals from Viking-associated contexts displaying diverse ancestries overlapping with both Scandinavian and pre-Viking British populations.^15^ Genomic analyses of two Viking Age mass graves in England (St. John’s College, Oxfordshire, and Ridgeway Hill, Dorset) support this complexity, revealing heterogeneous ancestry profiles that partially overlap with those of individuals traditionally classified as “Anglo-Saxon”.^15–18^ Despite these advances, the broader genomic legacy of Scandinavian settlement in England remains incompletely understood.^7,15,19,20^ Political and cultural entanglement between England and Scandinavia culminated in the 11^th^ century, when four Danish kings ruled England and the North Sea Empire unified Denmark and England under Cnut.^4^ These connections provided a direct pathway for the Norman claim to the English throne.^21–23^ Originating from Norse settlers in northern France during the 9^th^ century, the Normans developed a distinct identity blending Norse, Frankish, Breton, and Gallo-Roman influences, formalized with the establishment of the Duchy of Normandy in 911 CE.^4,24^ Norman influence in England predated the Conquest, particularly at Edward the Confessor’s court, and intensified after William I’s victory in 1066 CE, leading to profound political, administrative, and cultural transformations.^24–29^

Although the Norman Conquest is among the most intensively studied events in English history, its genetic impact remains unexplored. Previous archaeogenetic studies have focused on Roman, early medieval, and Viking-Age populations using both uniparental markers and genome-wide data,^1,7,15,30–32^ yet no genomic data from individuals directly associated with the Norman period are currently available. This absence limits our understanding of how the Conquest reshaped genetic diversity, particularly outside elite urban contexts. Archaeological and historical evidence suggests that while Norman influence is readily visible among aristocratic and urban populations, its effects on rural communities were more subtle and uneven.^26–29^

Here, we address this gap by generating and analyzing genome-wide data from 78 individuals, together with a newly compiled, consistently calibrated radiocarbon dataset for 98 individuals – largely overlapping with the genomic sample – from a rural cemetery in Surrey dating to the period surrounding the Norman Conquest. The cemetery, located in the Priory Orchard site in Godalming (POG) and associated with the Church of St. Peter and St. Paul, served the local population from the 9^th^ to the early 13^th^ century CE.^33,34^ Surrey occupies a strategically important position in southern England, with evidence of early Germanic settlement, subsequent Viking activity, and early Norman control following William I’s march toward London in 1066 CE.^35–39^ By examining genomic diversity in this rural population, we investigate how successive waves of continental and North Sea-mediated interactions, including the Norman invasion, shaped local genetic variation in the region. Our results provide the first genomic insight into communities living through the Norman Conquest and help refine models of medieval population history in England beyond traditional centers of power. In doing so, this study highlights the importance of rural contexts for understanding identity, integration, and demographic change during one of the most transformative periods in English history.

## RESULTS

### Archeological Site, Osteological, and Radiocarbon Dating Analyses

The Priory Orchard cemetery (Figure 1) in Godalming, Surrey, contains burial dating to the late Saxon, Viking, and early Norman periods. Used between the 9^th^ and 12^th^ centuries, it holds hundreds of burials reflecting a time of cultural transition.^33^ All individuals were buried supine and facing east, as is expected in a Christian cemetery of this date. The excavated site contained ∼300 interments. We randomly selected 78 individuals for paleogenomic analyses, based on the presence of preserved teeth or petrous portion of the temporal bone, and to ensure broad spatial representation across the cemetery.

**Figure 1:**
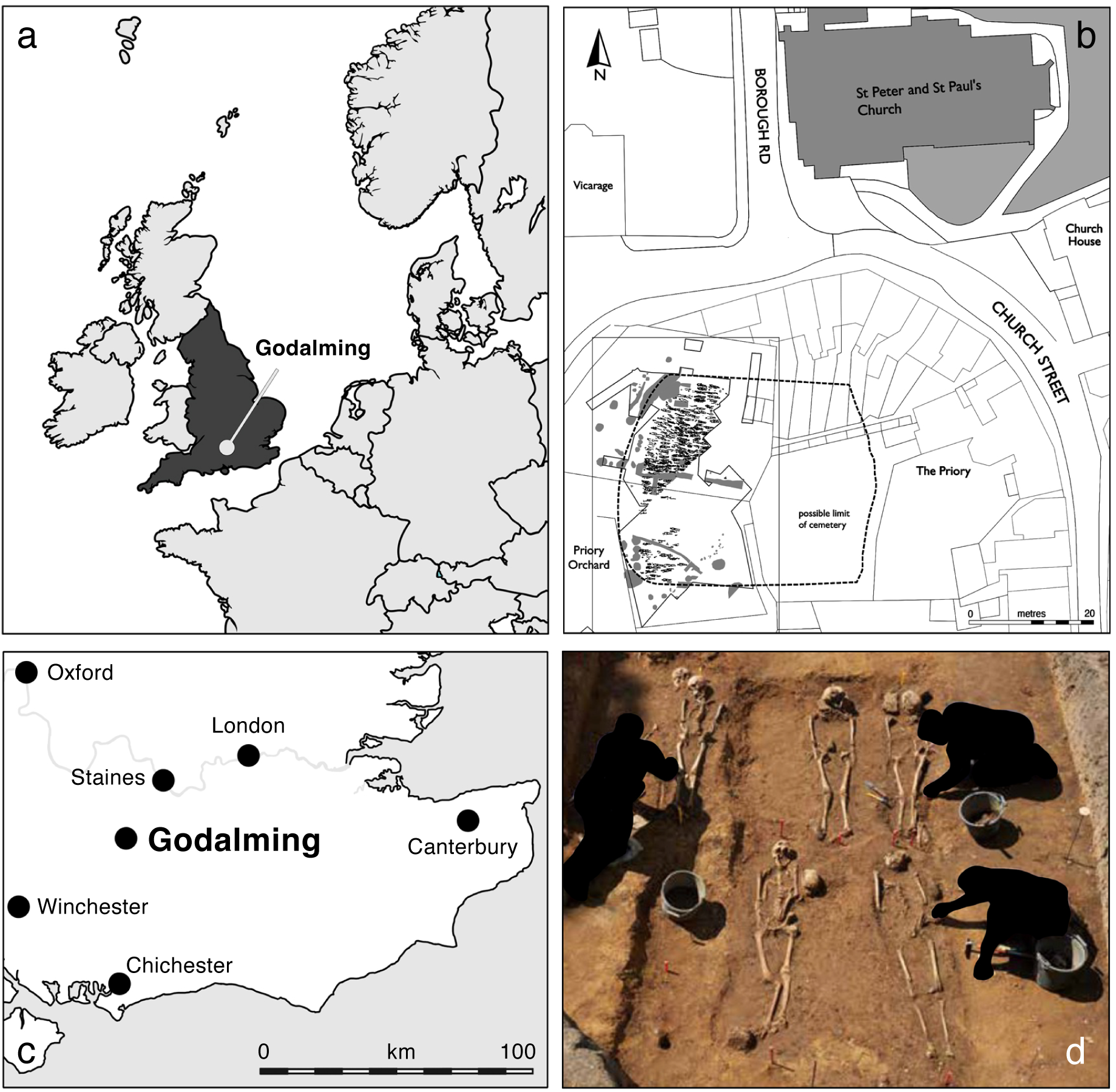
Location and excavation of the Priory Orchard of Godalming (POG) cemetery in Surrey, England. (a) Map showing the location of POG within northwestern Europe. (b) Archaeological site plan highlighting the cemetery’s position near St. Peter and St. Paul’s Church. (c) Map of southeastern England indicating the position of Godalming relative to major cities. (d) Archaeological excavation of multiple burials at the POG site.

Most of the individuals included in this genomic study along with a selection of additional skeletal samples were radiocarbon dated (103 dates across 98 individuals; Table S1). As part of a previous, independent study, collagen was extracted from post-cranial bones from 93 human individuals and underwent analysis via EA-IRMS for 8^13^C and 8^15^N determination as part of a larger study on Early Medieval diet in England using a modified Longin method.^12^ The leftover human collagen (n = 93) from EA-IRMS was then utilized for radiocarbon determination at the Oxford Radiocarbon Accelerator Unit, and combined with previously generated dates (n = 12) from the commercial excavation (see STAR Methods), for modelling in OxCal v.4.4, using mixed marine and terrestrial curves (Table S1).^33,40–43^ This radiocarbon framework provides a well-resolved chronological context for assessing genetic patterns across the Conquest period (Figure S1).

### Genomic Data and Authentication

The remains of 78 individuals were processed in the UCSC Human Paleogenomics Lab for DNA extraction. Single-stranded genomic libraries were prepared from DNA extracted from 76 teeth and four temporal bones, two of which were from selected individuals already sampled for teeth, and screened for ancient DNA (aDNA) using shotgun whole-genome sequencing. The raw pair-end sequencing data were filtered for endogenous DNA sequences by mapping reads to the human reference genome GRCh37 (hg19) and depth of coverage was estimated (Table S2). Of the 80 libraries, 41 had a mean coverage of less than 0.01x and were excluded from downstream analysis. Replicates derived from two specimens corresponding to a single individual did not contribute significantly to an increase in total yield beyond 0.01x. The remaining 39 libraries exhibited mean depth of coverage values ranging from 0.01x to 2.33x, with endogenous DNA proportions ranging from 0.34% to 89.66% (Table S2).

Genetic sex and mitochondrial and Y-chromosome haplogroups were assigned for individuals with sufficient coverage (Table S2; Document S1). Estimates of contamination based on the X chromosome in males and mitochondrial DNA in both sexes were consistently low (<2.5%; Table S2). Post-mortem deamination patterns were consistent with authentic aDNA, showing an excess of C-to-T substitutions at both ends of sequencing reads (Table S2).

### Genetic Ancestry Variation and Population Structure in 11^th^ Century Godalming

We selected 37 individuals who shared more than 10,000 single nucleotide polymorphisms (SNPs) with the 1240k dataset from the AADR v54.1 release.^44,45^ To avoid bias from close genetic relationships, we screened the cohort for first-degree and second-degree relatives and excluded one individual from each related pair (Table S3; Document S1). As a result, 19 individuals were excluded from downstream analyses, unless otherwise noted. Genomic data from the remaining 18 individuals were merged with published data from 1,616 ancient and modern European individuals from the AADR v54.1 release (Table S4).

As a first step to exploring the genetic diversity of the individuals buried in the POG site, we performed Principal Component Analysis (PCA), projecting the genetic variability of POG individuals – along with that of historical/ancient groups associated with multiple migratory waves into Southern England – onto the principal component space defined by a reference panel of present-day individuals representing genome-wide variation across the UK, Scandinavia, and parts of Western Europe (Figure 2a). The 18 unrelated POG individuals aligned with the distribution of UK samples along the first principal component (PC1); however, like UK Early Medieval individuals (5^th^-10^th^ centuries CE), POG individuals diverged from present-day UK populations along PC2.

**Figure 2:**
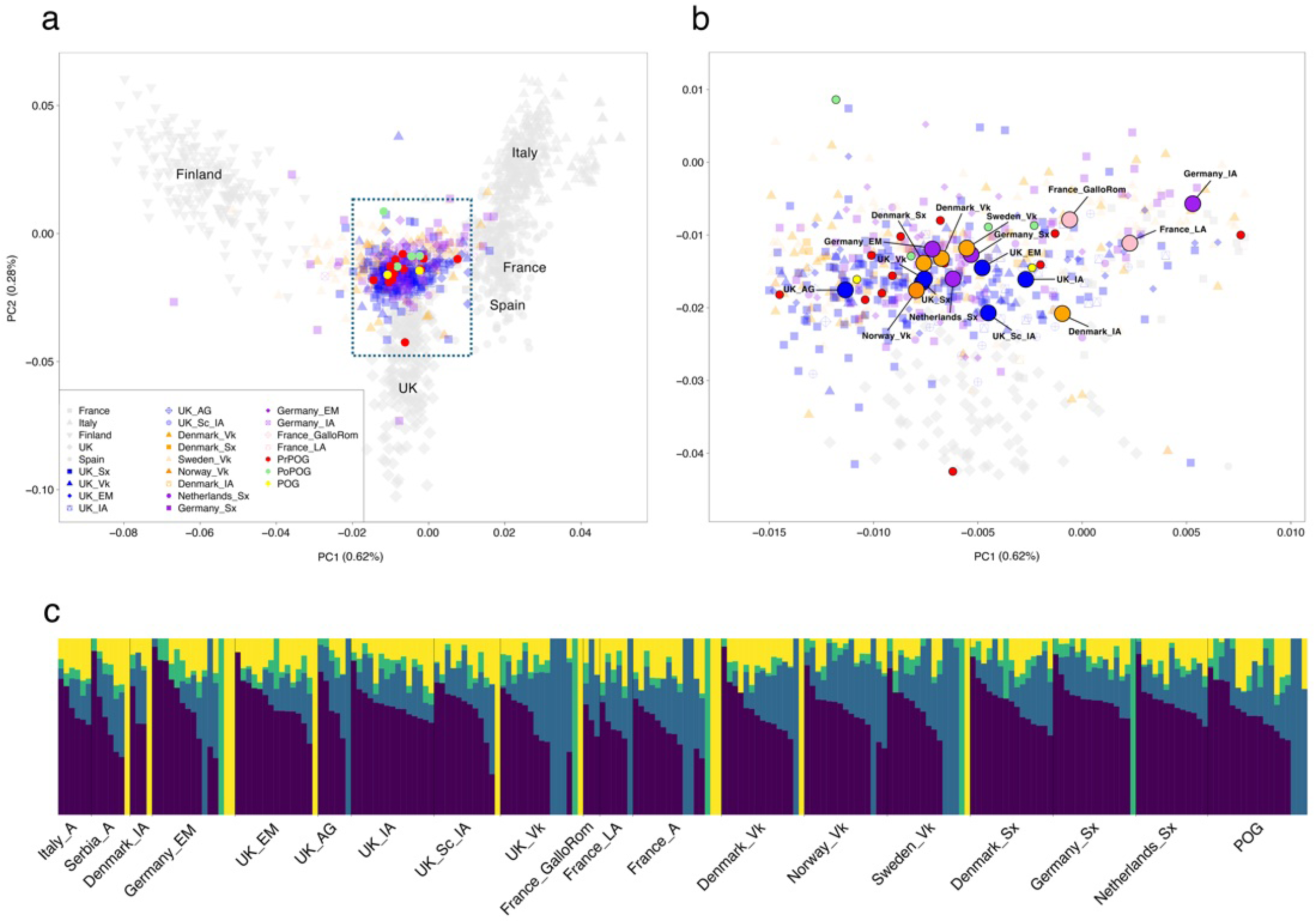
Principal component analysis and model-based clustering of Priory Orchard of Godalming (POG) individuals and comparative ancient populations. (a) Principal component analysis (PCA) based on present-day West Eurasian individuals from the AADR v.54.1 dataset. Ancient individuals were projected onto the resulting PC1-PC2 space. Gray symbols represent present-day individuals. (b) Zoom of the dashed region shown in panel (a). (c) Results of unsupervised ADMIXTURE analysis (K = 4) including POG and previously published ancient individuals. Abbreviations: Vk = Viking; Sx = Saxon; IA = Iron Age; EM = Early Medieval; AG = Anglian; GM = German; LA = Late Antiquity; A = Ancient. See also Figures S2—S4.

A comparison with roughly coeval individuals from Saxon, Viking, Early Medieval, and Late Antiquity groups across regions around the North Sea reveals that the POG individuals fall within the range of genetic variation of these ancient populations without forming clear clusters based on culture or period (Figure 2b). While the subtle differentiation among Medieval populations reflects the limited resolution of covariation-based methods such as PCA, it also underscores the high degree of interconnectivity among northern European populations during this period.^6,7,12,15,31,46–49^ These observations suggest that individuals living in England around the period of the Norman Conquest (10^th^-11^th^ centuries) exhibited diverse ancestry, but did not form clear population structure, consistent with previous findings for the early medieval period.^1,7^

A model-based clustering analysis was performed using both an extended cohort including 1,634 individuals and a subset of roughly contemporaneous individuals – 241 individual samples, with a maximum of 15 per population (Table S4) – to explore patterns of population structure in the POG population (Figures S2 and S3). The analysis with the 1,634-individual dataset was used to assess overall clustering stability and determine the optimal number of ancestral components across the full reference panel, whereas the 241-individual subset provided a balanced representation that facilitated clearer visualization of ancestry patterns. Results were broadly consistent between datasets. Initially, we tested unsupervised clustering across a range of cluster factors (K = 2-10; Figures S2 and S3) and determined that K = 4 minimized the cross-validation error (Figure S4). Similar to the PCA, this analysis, considering K = 4, revealed no marked differentiation between groups defined by cultural or temporal labels, although subtle differences in ancestry components were observed across individuals (Figure 2c).

To further elaborate on the genetic makeup of POG individuals, we employed a supervised clustering approach using Saxon-, French-, and Viking-associated samples (as defined in the AADR dataset^44^) as proxy source populations (Figure 3). We modeled the Saxon as a single source (yellow), and French samples (namely, France IA and France Late Antiquity/Gallo-Roman) were combined into another single source (blue), while Viking groups were partitioned according to their geographical origins (Denmark = purple, Norway = red, and Sweden = green). The analysis confirms that POG is characterized primarily by a Viking-related genetic component, with relatively smaller contributions from Saxon and French sources. Specifically, approximately 31% of POG’s genetic ancestry is associated with Swedish Vikings, followed by ∼21% associated with Danes, and ∼13% with Norwegian-like Vikings. According to this model-based clustering analysis, the Saxon component accounts for ∼29%, while only ∼6% of POG’s genetic ancestry is shared with the French group (blue).

**Figure 3:**
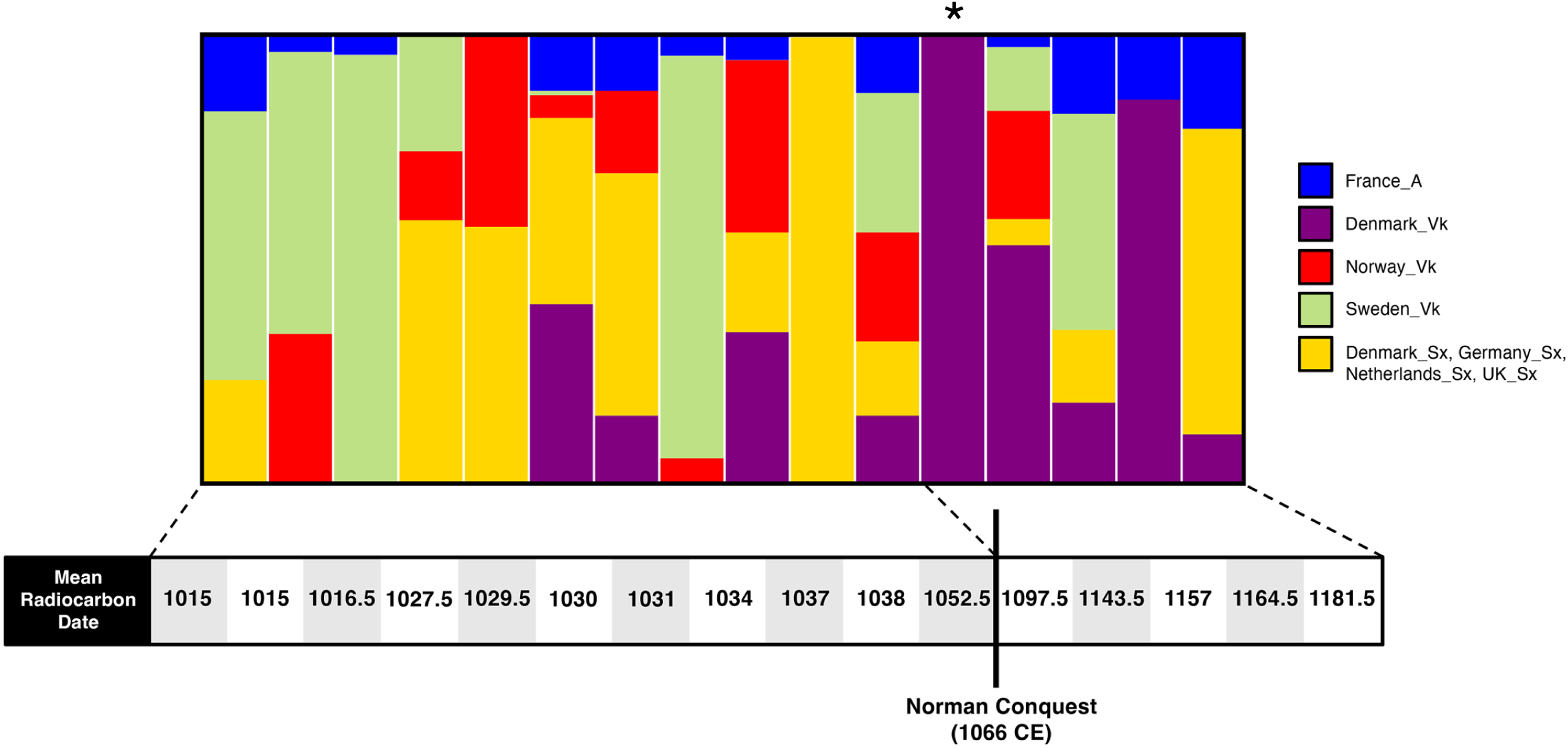
Temporal changes in inferred ancestry proportions from Viking and continental European sources. Results are based on a supervised model-based clustering analysis (see STAR Methods). Each vertical bar represents an individual ordered by mean radiocarbon date (Table S1). Colors indicate genetic affinities to reference populations: France (blue), Denmark Viking (purple), Norway Viking (red), Sweden Viking (light green), and Saxon-associated groups from Denmark, Germany, the Netherlands, and the UK (yellow). The vertical marker on the timeline indicates the Norman Conquest (1066 CE). Highlighted with an asterisk is individual POG15-3107, a young adult female who was buried with a linen smoother (Figure S5).

To investigate the potential genetic/demographic impact of individuals associated with the Norman Conquest, we stratified the POG samples based on their mean calibrated radiocarbon dates. We acknowledge that this approach overlooks the uncertainty inherent in radiocarbon dating; nonetheless, we adopted this simplified temporal stratification to investigate potential shifts in ancestry before and after the Conquest. This approach enabled us to assign 11 POG individuals with sufficient genetic data to the period before 1066 CE (PrPOG) and five to the period after (PoPOG). While no clear genetic differentiation is observed between Pre-Norman Conquest (PrPOG) and post-Norman Conquest POG individuals (PoPOG) in the PCA plot (Figure 2b), the supervised ADMIXTURE results (Figure 3) highlight the component associated with Danish Viking ancestry (purple) as relatively more pronounced in the genomes dated to the period after 1066 CE.

### Genetic Affinities with Ancient Populations from the North Sea and Continental European Groups

While PCA and ADMIXTURE offer useful insights into the genetic structure and diversity of POG and other Middle Ages groups, their ability to resolve fine-scale differences is limited when applied to these populations due to their low differentiation, as shown in Figure 2. To assess genetic affinities more sensitively between POG individuals and other medieval populations, we employed outgroup f3-statistics (f3out) as f3out(POG, *Y*; Yoruba). This statistic measures the amount of shared drift between POG and *Y*, a European population from the comparison panel, relative to the Yoruba (YRI) population from Nigeria, which serves as an outgroup. Calculated f3out values indicate shared genetic drift (i.e., genetic affinity) between POG and the population *Y* relative to the outgroup (Figure 4a).

**Figure 4:**
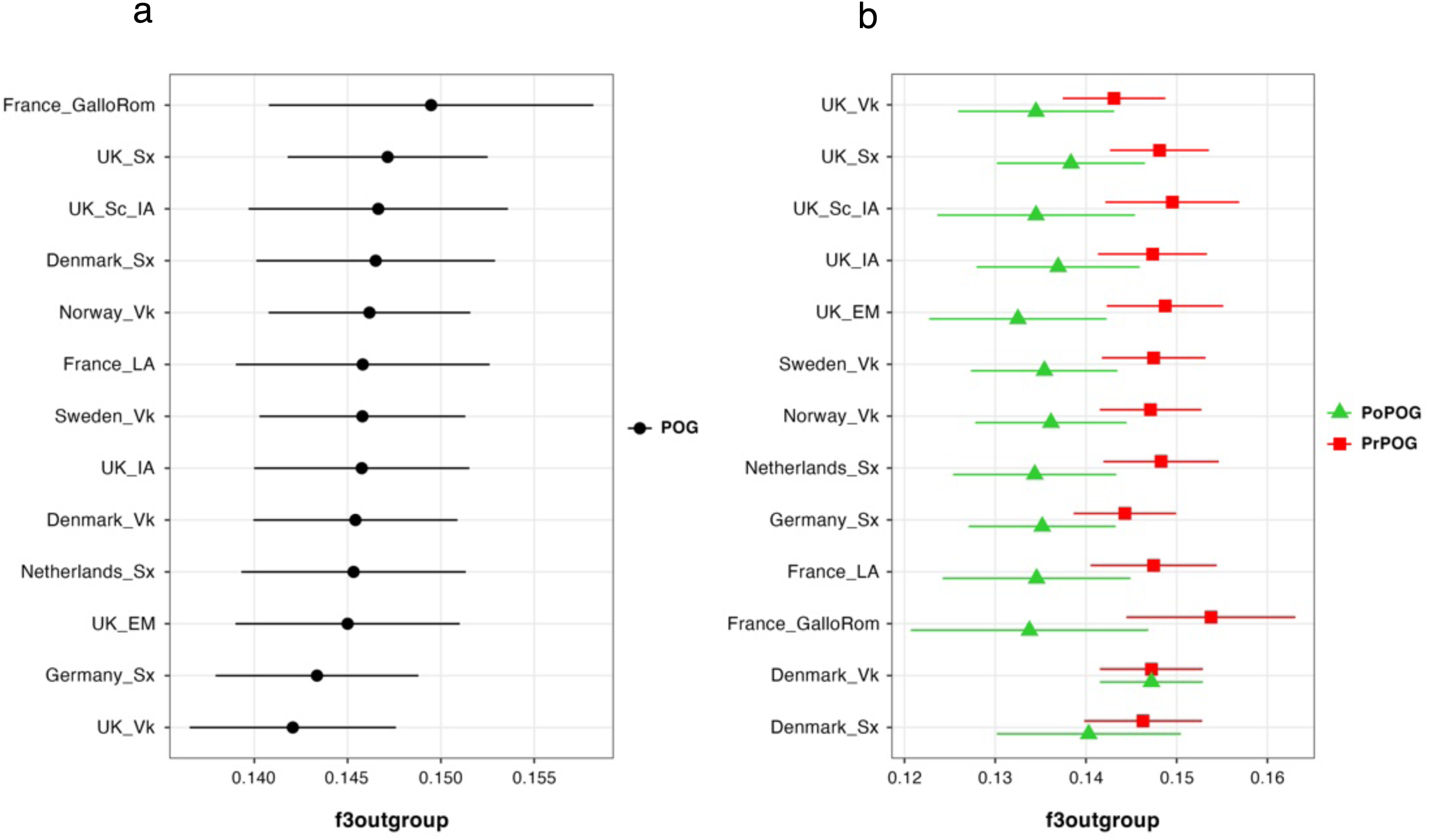
f3-outgroup statistics comparing genetic affinity among Priory Orchard of Godalming (POG) and ancient and modern European populations. (a) Outgroup f3 values showing shared genetic drift between the POG group and a range of European reference populations, including Viking Age Scandinavians, Saxon-associated groups, and continental Europan populations. (b) Comparative outgroup f3 statistics for PrePOG (POG samples dating pre-Conquest) and PoPOG (POG samples dating post-Conquest), illustrating differences in genetic affinity to the same reference populations. Points represent mean f3 values and horizontal bars denote standard errors.

Although the standard errors of the f3out estimates are relatively large, some trends can still be identified. The overall pattern in which POG shows strong drift-sharing with Atlantic Archipelago and Scandinavian populations suggests that POG’s ancestry was shaped predominantly by populations from the North Sea region. We note that although *France_GalloRom* shows the highest f3out point estimate (f3out = 0.1495), its standard error is the largest. This finding reflects, on the one hand, the limited sample size (n = 3) representing this group and, on the other, it suggests that POG has some genetic affinity to continental populations, particularly from France. Among the ‘Viking’ groups, which according to the model-based clustering analysis (Figure 3) are the major ancestry group in POG, *Norway_Vk* (f3out = 0.1462) shows the highest affinity (Figure 4a). The affinity with continental populations such as *Germany_Sx*, *Netherlands_Sx*, and *France_GalloRom* is perhaps less surprising when viewed within the context of the interconnected North Sea world, where longstanding trade, mobility, and cultural exchange – particularly through emporia like Quentovic, located near the mouth of the Canche River in northern France – fostered shared ancestry across Northwestern Europe.^50^

Stratifying the POG individuals into pre- (PrPOG) and post-Conquest (PoPOG) reveals a consistent trend toward lower f3out values in the latter (Figure 4b). While most comparisons produce overlapping estimates, considering the large standard error, the divergence is particularly notable when compared to Early Medieval individuals from the UK (*UK_EM*). Despite this general lack of significant differences in the f3out values for the time-stratified samples, there is a clear trend toward dilution of the genetic affinity, after 1066 CE, between POG and the tested Iron Age, Late Antiquity, and Medieval populations.

### Saxon and Viking Contributions to POG Ancestry Reveal Distinct Patterns

To further explore the relationships between POG and other roughly coeval populations, we used D-statistics. Specifically, we tested for asymmetry in shared drift between POG and a pair of comparison populations (Pop1 and Pop2), with YRI as the outgroup, considering the following model: D(YRI, POG; Pop1, Pop2). Despite the subtle genetic differentiation between the comparison populations, likely reflecting historical interactions and the brief time period for populations to accrue differences, some patterns emerge. Our results highlight complex interactions among Saxon-, Viking-, Early Medieval British-, and French-related ancestry sources (Figure 5a). Notably, POG shows excess allele sharing with *UK_Sx* compared to both *UK_Vk* (D = −0.0044, Z = −3.654) and *UK_EM* (D = −0.0052, Z = −3.456), suggesting an affinity between POG and roughly contemporaneous and geographically proximate populations, to the exclusion of Early Medieval groups and others with origin outside the UK (*UK_Vk*). However, comparisons involving *Sweden_Vk* suggest greater affinity with POG relative to *UK_Vk* (D = −0.0040, Z = −3.243), *Denmark_Vk* (D = −0.0033, Z = −2.986), and even *UK_Sx* (D = −0.0032, Z = −2.776) (Figure 5a). Overall, these results are consistent with our previous observations, implying that POG exhibits a closer genetic affinity with Scandinavian and northern European Viking-related populations than with continental European groups.

**Figure 5:**
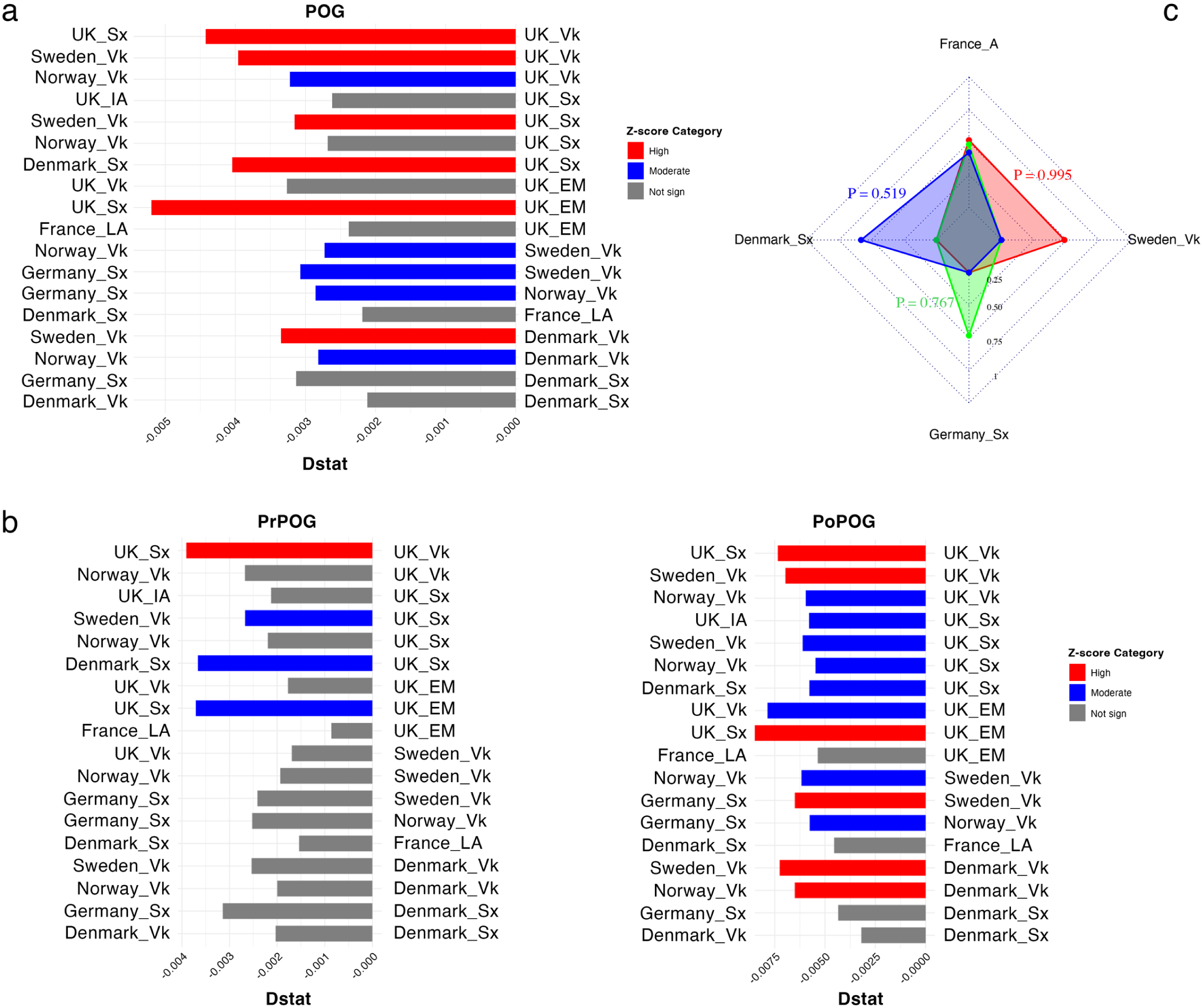
D-statistics and qpAdm modeling results. (a) Results of the D-statistic tests D(YRI, right; left, POG), including all POG individuals suitable for analysis. Abbreviations: Vk = Viking; Sx = Saxon; LA = Late Antiquity. (b) Results of the D-statistic tests D(YRI, right; left, PrPOG) and D(YRI, right; left, PoPOG). (c) Significant and biologically plausible qpAdm models using YRI, Spanish, ALG, and Italy_A as outgroups.

As with the f3out analysis, stratifying the POG samples into PrPOG and PoPOG based on their chronological context allowed us to assess potential shifts associated with the arrival of the Normans in the region (Figure 5b). For the PrPOG group, four D-statistic values were statistically significant, demonstrating increased allele sharing with UK Saxon and more so with Scandinavian populations relative to other groups. We note that *UK_Vk* comprises individuals from massacre sites and who do not have a unique origin outside the UK.^15–18^ The tests considering another Viking-related population (i.e., *Sweden_Vk*), as well as groups of Germanic-speaking people settled in Britain (*UK_Sx*) and from Denmark (*Denmark_Sx*), yielded trend-significant Z-scores exceeding 2, but that did not reach strong significance (i.e., Z scores ≥ 3), highlighting the slightly higher affinity of PrPOG with Scandinavian populations than with *UK_Sx*. Considering PoPOG, most of the D-statistics analyses suggest that Scandinavians show greater allele sharing than other Continental and Atlantic Archipelago groups. The sole exception appears to be the comparison involving *Germany_Sx* vs. *Sweden_Vk*, which suggests higher affinity of POG with the former (D = −0.0065; Z-score = −2.571). Overall, the partially overlapping results between PrPOG and PoPOG highlight how the region’s genetic homogeneity may limit the resolution of population differentiation, as the different human groups share subtle and closely related genomic signatures.

As a complementary approach to assess the complexity of ancestral contributions to POG, we performed qpWave analyses using multiple combinations of reference populations. These analyses, together with the corresponding qpAdm models (see below), were conducted using the pooled POG sample only (i.e., without subdivision into PrPOG and PoPOG) to minimize overfitting and spurious inference. Across most tested configurations, qpWave supported a single distinguishable ancestry stream for POG relative to the reference populations considered (Table S5). Specifically, qpWave analyses using Yoruba (*YRI*) as an outgroup and ancient French individuals (*France_Late Antique*, *France_GalloRom*, and *France_IA*; hereafter *France_A*) as candidate sources, together with *Italy* or *Spain* as right populations, yielded p-values of 0.149 and 0.090, respectively, indicating no rejection of a one-stream model. Similarly, when Mozabites from Algeria (*ALG*) or ancient Serbians (*Serbia_A*) were included among the right populations, the corresponding p-values were higher (p = 0.748 and p = 0.235). Consistent results were obtained when populations with high genetic affinity to POG, such as *Germany_Sx* and *Denmark_Sx*, were used as right populations, with p-values generally exceeding 0.05. In comparisons involving Scandinavian Viking Age individuals (*Sweden_Vk*), qpWave revealed significant allele-sharing asymmetries between POG and southern or southeastern European populations (e.g., Italy and Spain, p = 0.0029 and p = 0.0022, respectively; *Italy_A* and *Serbia_A*, p = 0.0308 and p = 0.0195), consistent with greater genetic affinity of POG to northern European groups. The increased allele sharing with Scandinavian Viking Age individuals reflects shared northern European genetic ancestry rather than evidence of direct Viking ancestry or migration into POG. Nevertheless, these comparisons did not support *Sweden_Vk* as a distinct additional ancestry stream, as the one-stream model could not be rejected when *Sweden_Vk* was included among the reference populations. The occasional rejection of the one-stream model, particularly in comparisons involving temporally or geographically distant ancient populations, does not imply the absence of admixture, but likely reflects limited power to distinguish multiple closely related ancestry sources with substantial shared genetic background. Notably, most non-rejected models (p ≥ 0.05) involve continental Saxon and Frankish groups (Table S5), consistent with POG displaying a genetically coherent profile closely aligned with continental Saxon and Frankish populations; any additional ancestry components, if present, are likely closely related and not readily distinguishable.

Given qpWave results indicating that POG is best described by a single ancestry stream relative to the tested reference populations, we next applied qpAdm to assess whether this genetic profile could be parsimoniously represented as a mixture of two closely related European ancestry sources and to estimate their relative contributions. In this context, the qpWave one-stream result reflects limited power to distinguish among highly correlated ancestral sources with substantial shared genetic background, whereas qpAdm provides a flexible framework for approximating such continuous variation using a small number of reference populations. Thus, the two-source qpAdm models should be interpreted as simplified representations of ancestry along a genetic cline rather than evidence for discrete admixture events.

Using rotating outgroups including Yoruba (*YRI*), *Italy*, *Spain*, and Algeria (*ALG*), we observed that POG was consistently well fit by two-way models combining northern/central and western European reference populations, with p-values generally exceeding 0.05, indicating that these models cannot be rejected. A representative model estimates POG ancestry as 51.6% *France_A* and 48.4% *Sweden_Vk* (p = 0.995), while an alternative model using *Germany_Sx* and *France_A* yields comparable proportions (48.3% and 51.7%, respectively; p = 0.767). Similar statistical fits were obtained when *Denmark_Sx* was substituted for *Germany_Sx* or *Sweden_Vk*, further underscoring the non-uniqueness of the inferred source populations (Table S6; Figure 5c).

In several cases, models produced small negative coefficients despite high p-values; while ancestry proportions cannot be negative in a biological sense, such estimates commonly arise in qpAdm when closely related source populations share substantial genetic ancestry and are difficult to distinguish statistically^51^, rather than indicating true absence of contribution. Taken together, these results place POG along a North-West European genetic cline and indicate that its ancestry can be adequately described by mixtures of broadly northern/central and western European-related components, without evidence for a uniquely Scandinavian or temporally specific admixture event.

## DISCUSSION

Here, we reconstruct the population history of early medieval Surrey using genomic data from 78 individuals buried at Priory Orchard and dated from the 9^th^ to the early 12^th^ centuries CE. We evaluate the genetic impact of the Norman Conquest on this rural population and place these individuals within the broader demographic landscape of the Middle Ages. In the sections below, we show that the genetic makeup of the POG population reflects substantial Viking-associated ancestry alongside important contributions from continental Europe. We then examine the genetic legacy of the Norman Conquest in Surrey and discuss the broader implications of our findings for understanding population structure and mobility in medieval Europe. The broader historical and archaeological framework relevant to these findings is discussed in Document S1.

### Scandinavian Influence in Southern England

POG shows notable genetic affinities with Scandinavian populations from the Viking Age, particularly from Denmark and Sweden. Although Viking activity in Surrey has not been widely discussed in the literature, there is evidence that the region was not just an occasional target of Viking raids. According to the Anglo-Saxon Chronicle,^52^ Viking armies moved south across the Thames and through Surrey in the 9^th^ century culminating in the battle of Acleah, which had previously been identified as Ockley, in Surrey (close to Godalming), but which most scholars agree is more likely to have been Oakley, in Hampshire.^53–55^ The construction of a defensive fortification in Eashing (only 2 km distant from Godalming) around 885 CE^38,39^ suggests that Viking raids were a significant threat and this is confirmed by the Battle of Farnham (893 CE)^37^ (12 km from Godalming and 10 km from Eashing), where an army led by Edward, son of King Alfred, was victorious over the Danes. That these incursions translated into a significant permanent Viking presence is suggested at Guildford (6 km from Godalming), which replaced Eashing as the area’s defensive center (burh) for Wessex in the 10^th^ century CE.^56^ Its Late Saxon plan can be defined with some confidence, with a single main street with side streets leading to the defenses. A number of the side streets have the suffix ‘gate’ (*Tun Gate* and *Swan Gate*, for example, as named on a map of 1739).^57,58^ This can only be the result of Scandinavian influence since the term is derived from Old Norse and in England this usage is otherwise only known from the north (e.g., York).^59^ Some other, generally more distant, indicators of a Scandinavian presence in Surrey can also be mentioned. The wealthy and important Abbey at Chertsey (24 km from Godalming) was twice sacked by Vikings in 871 CE and the early 10^th^ century CE.^60^ It is tempting to speculate that a Viking sword found in a former channel of the Thames at Chertsey,^61^ belonged to one of these raiders. Whatever the truth of that, it forms one of a range of objects with Scandinavian links from the reaches of the Thames in this area, part of an efflorescence of ritual (votive and pagan) deposition of artefacts in the river at this period.^62^ To this may be added, from various parts of Surrey, a small number of Scandinavian style finds recorded on the Portable Antiquities Scheme dating primarily to the 11^th^ century^63–65^ and occasional finds from excavations, such as a pre-Conquest bone ice skate from Reigate (30 km from Godalming) of a type otherwise only identified in England in the Danelaw.^66^

These observations, together with our findings reinforce the view that the Scandinavian presence in Surrey was not fleeting. The significant Viking-associated genetic ancestry in POG highlights the lasting impact of Scandinavian settlements on the region, especially in the 11^th^ century under Danish rule. Additionally, the contribution of Swedish Viking-like ancestry to POG underscores the expansive reach of North Sea networks. Supporting this, the D-statistics revealed increased allele sharing between POG and Scandinavian populations, and the combination of qpWave and qpAdm analyses provides additional evidence for a significant Viking-Age Scandinavian contribution to the POG population. These findings reflect the interconnectedness of the Viking world and the enduring genetic and cultural legacies they left in southern England.

Consistent with these findings, POG15-3107, a young adult female individual, was buried with a linen smoother (Figure S5), one of only three items (the other two were spindle whorls) deliberately placed with burials in the cemetery, likely to indicate a person of exceptional status. These objects were used in the finishing and laundering of textiles and garments and have been discovered at archaeological sites in Britain dating from the 8^th^ century into the Medieval period,^67^ typically associated with female individuals and documented across Viking territories such as Anglo-Scandinavian York and Winchester.^68^ In Scandinavia, linen smoothers are relatively common finds in graves with females individuals during the Viking Age and are frequently interpreted as symbols of high status.^33,69^ Indeed, a prominent example of a linen smoother found in a funerary context in England is from the richly furnished Viking burials of the 10^th^ century at Cumwhitton in Cumbria.^70^ Notably, POG15-3107 also displays substantial proportion of Scandinavian-related ancestry among the analyzed individuals in the individual-level, supervised ADMIXTURE analysis (Figure 3), providing a striking correspondence between the genetic data and the culturally diagnostic grave goods.

### Genetic Evidence of Continental Interactions in Surrey

As previously noted, the POG cemetery likely served a population drawn from the large area known as the Godalming Hundred. That entity is likely to have emerged prior to the earliest burials at POG and it seems it was a subdivision of a larger *regio* (sub-kingdom) of the Godhelminghas that occupied all of south-west Surrey.^34^ Notably, no Early Saxon burial sites are known from this territory, in contrasts to their prevalence over much of the rest of Surrey.^71^ This absence suggests that there was no significant Saxon cultural, and presumably population, presence in the area until after the late 7^th^ century, when this type of burial rite largely ceased. Most of the area occupied by Godalming Hundred belonged to the Weald, an area where there was a significant expansion of settlement as it was converted from extensive pasture to more intensive agricultural use in the period between the 8^th^ and 12^th^ centuries.^34^

Across multiple allele-sharing-based analyses, including D-statistics, we observe a persistent but moderate Saxon-related influence, with tests involving *UK_Sx* showing significant but comparatively weaker signals than those associated with Scandinavian-related populations. Importantly, the genetic signatures support a long-term history of demographic interactions across the North Sea, resulting in a population shaped by both local Saxon-related ancestry and continental inputs. At a broader scale, populations across England experienced near-exponential growth, rebounding from a low point during the Early Saxon period.^33^ It is therefore plausible that migrants arriving after this Early Saxon demographic bottleneck, rather than the initial Saxon settlers, contributed substantially to the genetic composition observed at POG. This pattern likely reflects early migrations from the Jutland peninsula and northern Germany during the 5^th^ and 6^th^ centuries and may also capture later influences from Viking groups with shared Saxon heritage, as suggested by the consistently higher allele sharing between POG and *Danish_Sx*, compared with *UK_Sx*. Rather than indicating a single episode of population replacement, these patterns point to a population in southern England shaped by overlapping continental and North Sea connections and sustained population exchanges over several centuries.

### The Norman Conquest and Its Genetic Legacy in Surrey

The presence of French-associated ancestry in some of the analyses aligns with historical narratives of migration, trade, and demographic exchange during the early and central medieval periods in southern England. For instance, qpAdm modeling estimates that approximately 51.6% of POG ancestry can be modeled using an ancient French population as a proxy source. This scenario likely reflects – but is not solely attributable to – shifts in elite structures following the Norman Conquest of 1066 CE, which involved the arrival of individuals from across northwestern Europe, including Normans, French, Flemish, and Bretons.^72–75^ It may also support historical suggestions of a Breton presence at the West Saxon court prior to 1066, which has until recently been ephemeral in historical and archaeological studies.^76,77^

Beginning in the late 8^th^ century, Scandinavians – primarily from Denmark, Norway, and Sweden – launched raids along the northern coast of the West Frankish Kingdom (roughly corresponding to modern France).^4,78^ These raids targeted wealthy monasteries, villages, and cities along the rivers. Over time, some Vikings began to winter in these coastal and riverine areas, establishing semi-permanent camps and later, settlements in the region of Normandy.^4,78,79^ By the 11^th^ century, these settlers – the Normans – had expanded across Normandy and developed a unique identity, gaining a reputation as formidable warriors and skilled organizers. As a result, they became sought-after mercenaries across Europe.^4,5,80^

In the first half of 1066 CE, between 7,000 and 12,000 Norman soldiers were deployed to England, arriving aboard a fleet of approximately 600 to 700 ships.^81–83^ Under William’s leadership, they landed first at Pevensey Bay, setting the stage for the Battle of Hastings on 14 October 1066. This decisive battle, fought between William’s Norman-French forces and the English army led by Harold Godwinson, marked the beginning of Norman dominance in England. Following their victory, the Normans quickly secured control over the entire country.^80^ Their advance into Surrey began soon after their arrival in Britain, with Normans extending their control over southern England even before William claimed the throne. Their first significant stop was Dorking, just a few kilometers east of Godalming. There, William’s army was reinforced by troops from newly conquered territories in southern England before continuing their march northward toward London.^36^

The French-associated genetic ancestry we detect in POG aligns with this historical context, where the establishment of Norman aristocracy and the integration of their descendants into English society facilitated the flow of continental genes. Signals of affinity with French populations emerge primarily in specific admixture modeling frameworks, particularly qpAdm and, to a lesser extent, f-outgroup statistics, rather than uniformly across all analytical approaches. While multiple continental sources contribute to the genetic profile of POG, these results suggest that French-related ancestry represents one plausible component within a broader spectrum of northern European influences, rather than a singular or dominant source. Moreover, the French-associated contribution might not be exclusively attributable to the Norman Conquest itself; intensified cross-Channel exchanges in commerce, culture, and population movement both before and during this period likely facilitated the gradual incorporation of French-related genetic elements into the population of southern England.^84^

But did the Norman Conquest significantly reshape the genetic landscape of rural southern England? A comparison between PrPOG and PoPOG individuals reveals only subtle shifts in ancestry and genetic affinity that do not reach statistical significance. These observations align with historical and archaeological sources that challenge the notion of abrupt cultural and demographic change following the events of 1066.^26–29^ Evidence highlights more drastic changes to elite and political structures, as well as land ownership, particularly in the urban centers. However, most aspects of rural life remained relatively undisturbed. This picture of continuity rather than disruption is particularly evident in southern England.^27,28^ Our study thus offers an unprecedented opportunity to re-examine continuity and change in the aftermath of the Conquest, showing that continuity reflects sustained admixture with Northern European populations and broader regional population exchanges rather than a stable isolated population or a sudden Norman genetic influx.

Despite the general lack of significant differences in the f3out values before and after the Conquest, there is a clear trend toward dilution of the genetic affinity between POG and the tested populations after 1066 CE. Several factors could account for this dilution. First, the individuals arriving in Surrey around 1066 CE may have carried ancestry from a group that is either not well-represented in our chosen comparison panel or genetically more heterogeneous than the tested groups. This novel input could either dilute the shared drift between POG and the tested groups or increase ancestry variance within PoPOG, thus consistently lowering f3out values. Alternatively, a drop in f3out could also result if PrPOG individuals were more genetically homogeneous than PoPOG. Lastly, local genetic drift could have contributed to lower f3out values after 1066 CE, though this is unlikely given the relatively short elapsed time.

Beyond these nuanced shifts in f3out values putatively associated with the Conquest, no major changes are observed in the genetic pool of POG. These findings contribute to long-standing scholarly debates about the nature and impact of the Norman Conquest in England, and prior Breton presences in the region.^26–29,76,77,80,80,82^ Our genomic data from POG show only minor perturbations in the local gene pool, resonating with narratives that emphasize continuity rather than deep transformations in rural England during the mid-11^th^ century. Specifically, our data support the view that the Norman Conquest did not trigger a widespread demographic turnover in rural communities of the south. While the genetic data suggest continuity, we note that the archaeological record at POG may point to a nuanced cultural shift reflecting continental influences.^85^

Taken together, our findings reveal that the legacy of the Norman Conquest in rural southern England was more complex than simple population replacement. While cultural and political change is clearly reflected in the archaeological record, the genetic data from POG instead point to substantial demographic continuity, shaped by long-standing cross-Channel interactions rather than abrupt disruption. Moreover, this case study illustrates the value of integrating genomic and archaeological evidence to resolve how major historical events translated into population-level outcomes.

### Implications for Medieval Population Dynamics

The genetic heterogeneity observed in POG individuals highlights the complexity of medieval population dynamics. The convergence of Scandinavian, Frankish, and English influences shaped the region’s diverse ancestry and culture. Our findings illustrate southern England as a vibrant crossroads of migration and cultural exchange during the Medieval period, revealing a region shaped by the interaction of diverse peoples and influences. The profound impact of Viking migrations is evident in the genetic signatures they left behind, marking centuries of raids, settlements, and integration that began in the late 8^th^ century. Viking groups, particularly from Denmark, established significant footholds across England, most notably in the Danelaw region, where their influence extended beyond genetics to include language, culture, and governance.^86,87^ Undoubtedly, the Norman Conquest of 1066 added another transformative layer to this already complex genetic mosaic. Norman rulers introduced continental practices, established a feudal system,^29^ and brought their own genetic diversity. The influx of Norman aristocrats, soldiers, and their families from across France, Brittany, and Flanders may have further diversified the genetic landscape of England, integrating with the existing heritages.

The genetic differences observed in pre- and post-Conquest POG suggest not only the arrival of new populations but also the ongoing integration and exchange between established and incoming groups. The Normans’ influence in England was already palpable before their conquest, as they had secured a strong foothold in Britain long before their decisive victory at Hastings.^88^ As a people of Scandinavian descent who had settled in the region of Normandy, the Normans played a key role in shaping the political and cultural landscape of both France and England, and their genetic legacy can be effectively recovered in the genomic landscape of POG, as our results show. Together, the genetic and historical patterns observed in POG individuals from Surrey portray a dynamic and evolving demographic landscape. Southern England during the medieval period was far from static; it was a nexus of cultural and genetic exchange. Different peoples and traditions converged and blended, leaving an indelible mark on the history and people of the region.

To gain a more comprehensive understanding of these historical dynamics, future research should prioritize the generation of high-coverage ancient genomes and the refinement of admixture dating techniques. Such advancements would allow for more precise reconstructions of genetic timelines, offering deeper insights into when and under what circumstances different populations contributed to the genetic makeup of medieval southern Britain. Integrating genetic data with information from archaeological artifacts, burial practices, and isotopic analyses will provide a richer, more nuanced picture of human life during this period. These combined approaches will help link genomic evidence to the lived experiences of past communities, offering a fuller understanding of their history. Our study underscores the remarkable potential of ancient DNA to illuminate the complex interplay of migration, culture, and genetics in shaping human history. By tracing the genomic legacies of Viking incursions and the Norman Conquest, this study advances our understanding of how these migrations influenced medieval communities and left a lasting imprint on the genetic heritage of modern populations in England.

## STAR⍰Methods

### Archaeological Context and Human Remains

The human skeletal material examined in this study was part of the Priory Orchard osteological collection, currently housed in the Department of Anthropology, University College London. The Priory Orchard osteological collection includes all the remains excavated in 2012—2015 from the Priory Orchard cemetery. The site is located within the Godalming Conservation Area and an overlapping Area of High Archaeological Potential. In 2007, archaeological human remains were uncovered during the excavation of a soakaway in the northeastern section of the site. In November 2012, as a requirement of planning approval, a trial trench evaluation was carried out and revealed that the eastern part of the site contained a densely populated Christian cemetery.^33^ The initial phase of systematic excavation, conducted in spring of 2014, covered the area where planned buildings were to be constructed over the presumed burial ground site. This phase defined the burial ground’s north-west boundary, uncovering 73 in situ inhumations along with a considerable amount of disarticulated bone, all of which were documented and removed.^33^ In early 2015, a phase exposed the west and south-west limits of the burial ground, with an additional 225 inhumations recorded and removed. The inhumations were systematically evaluated to estimate age-at-death, sex, and evidence of pathologies, using standard osteology methods. The articulated skeletons included 23 children and 275 adults, of which 75 were identified, based on osteological characteristics, as probable females and 126 as probable males^89–91^.

According to our extensive radiocarbon dating survey (Table S1; Figure S1), the cemetery was in use in from the 9^th^—13^th^ centuries CE. As a rare example of a large and mostly undisturbed rural cemetery of this age, the remains have exceptional research value, and their preservation for future research was agreed with the relevant stakeholders, including representatives of the local town and county council, the Godalming Museum, the archaeology and osteology research teams, the university accepting custody of the remains, and the local church. In line with this agreement, 76 teeth and four temporal bones (two of the temporal bones originated from an individual for whom a tooth was also analyzed) were chosen for the genomic analyses (and separate isotope study), which were performed following an ethically responsible approach to human remains.^92^ These 78 individuals include four children (age < 18), 28 young adults (most likely 18-29), 32 middle age adults (most likely 30-45), five older adults (most likely >45), seven adults likely middle age or older, and two adults of indetermined age. The adults included 26 probable females and 42 probable males, with the rest of indeterminate sex.

### Radiocarbon Dating of the Human Remains

Twelve bone and tooth samples were sent for radiocarbon determination during the initial phases of work by Surrey County Archaeological Unit. Analysis and initial calibration using IntCal13 was conducted by Beta Analytic via accelerator mass spectrometry, quoting 1 sigma errors, and 0^13^C and 0^15^N values were also analyzed by Beta Analytic using isotope ratio mass spectrometry (IRMS) in a similar fashion to our own isotopic analyses reported below.^33,93^

An additional 93 human bones (63 rib, 21 long bones, eight unidentified bone fragments, one vertebral fragment) and 16 faunal bones were analyzed for 0^13^C and 0^15^N as part of a separate study, with the remaining collagen then later used for additional radiocarbon dating and modelling here.^12^ For the human burials, ribs were preferentially sampled from humans, with other long bones chosen if ribs were unavailable, as ribs have a faster turnover rate and therefore should represent dietary carbon intake from the decade before death.^94,95^ Collagen was extracted from bone samples following a modified Longin method at the Dorothy Garrod Laboratory, McDonald Institute for Archaeological Research, University of Cambridge.^96,97^ Aliquots of collagen were run in triplicate using an automated elemental analyzer coupled in continuous-flow mode to an isotope-ratio-monitoring mass-spectrometer in the Godwin Laboratory, Department of Earth Sciences, University of Cambridge. Stable isotope values of bone collagen and dentine are reported relative to internationally defined scales, VPDB (δ^13^C) and AIR (δ^15^N). Analytical error (1σ) for all collagen samples was ±0.20‰. As per Leggett et al.^96^, the standards used are IAEA standard of caffeine for carbon and nitrogen (δ^15^N +1.0/1.1‰, δ^13^C −27.5‰); in-house standards of nylon (δ^15^N −3.14‰, δ^13^C −26.55‰), alanine (δ^15^N −1.4‰, δ^13^C −26.9‰), “new” alanine (δ^15^N −1.22‰, δ^13^C −23.88‰), protein 2 (δ^15^N +6.0‰, δ^13^C −26.9/-27.0‰) and EMC (Elemental Microanalysis caffeine, δ^15^N −2.5/-2.6‰, δ^13^C −35.8/-35.9‰) for carbon, nitrogen and atomic C/N ratios. All 109 POG remains samples successfully passed established collagen quality criteria at this stage.^12,98^ Full details on laboratory methods and protocols used for bone preparation and collagen extraction, are detailed in Leggett et al.^99^

Ninety-two human bone collagen samples with sufficient product left after EA-IRMS measurement were then sent for radiocarbon determination at the Oxford Radiocarbon Accelerator Unit (ORAU) using established methods.^40^ Three collagen samples failed (POG1033, POG1100, and POG3179), with additional bone fragments from skeletons 1033 and 3264 supplied as replacement samples. This gave 91 new radiocarbon dates, with five individuals having dates from both Beta and Oxford (POG1019, POG1023, POG3107, POG3258, POG3281). In total this gives 103 dates (12 Beta, 91 Oxford) across 98 individuals. Only five of the individuals with genomic data are undated (POG1100, POG3179, POG3339, POG3340 and POG3370a).

(Re-)calibration and Bayesian modelling of all dates were done using the IntCal20 and Marine20 radiocarbon curves in OxCal v.4.4.4.^41–43^ Using methods from previous Viking Age radiocarbon projects,^100,101^ the stable isotope results were used to calculate the percentage marine intake based on a terrestrial end member value taken from the cows at the site = ^13^C: −22.9‰;^96^ and a marine end member value was chosen from medieval Southampton as it was the most extreme ^13^C across the closest near-contemporary sites with fish isotopic data available: ^13^C: −11.5‰ (Southampton, Lower High Street - Cod).^96,102^ The equation used to calculate the fraction of marine dietary protein (f_m_) for any given individual *n* is:

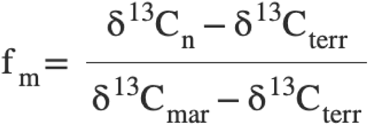

where ^13^C_n_ is the stable isotopic measurement of the individual’s sample *n*, ^13^C_terr_ is the fully terrestrial end member, and ^13^C_mar_ is the fully marine end member. A generalized uncertainty of 10% was included in the f_m_ estimates in OxCal at the expense of precision to account for propagation error.^100^ Estimated percentage marine protein dietary intakes from this model ranged from 7.7-30.6% across POG. It must be noted that this is a very simplistic measure using a linear mixing model and should not be taken as a true indicator of marine dietary intake in the population - however, it is useful for the purposes of accounting for marine reservoir effects in date (re-)calibration.^100,103^ A weighted mean ΔR value of = −166 was calculated using the ^14^Chrono Marine Reservoir Database (http://calib.org/marine/) drawing on the 10 nearest ΔR values to the site,^104–106^ which also conveniently corresponds to a suitable ΔR value for those with possible childhood origins in northern France, as this area of England lacks any suitable localized ΔR values.^100,101,103,106^

All data and code used for marine reservoir calculations and date calibration, and results are provided in the supplemental material online (Table S1; Data S1).

### Chronological Classification of POG Individuals

All POG individuals were initially managed as a single group for the analyses. However, due to the wide radiocarbon age range of the POG individuals and historical records noting the large-scale arrival of foreign armies in the area in 1066 CE, we divided them into two groups for some analyses. Those with an average radiocarbon date - in particular the midpoint between the lower confidence interval (CI) and the upper CI of the radiocarbon dating estimate - earlier than 1066 CE were categorized as “PrPOG,” while those with an average date later than 1066 CE were categorized as “PoPOG.”

### Ancient DNA Extraction, Library Preparation, and Sequencing

We analyzed 76 teeth and four petrous bone samples collected from the POG cemetery. The lab work was conducted in dedicated clean-room facilities at the Paleogenomics Laboratory, University of California, Santa Cruz (UCSC-PGL; Santa Cruz, CA, USA). Teeth were mechanically brushed to remove soil traces and soaked in 3% Sodium Hypochlorite solution to clean the outer surfaces of both crown and root. After rinsing in molecular biology-grade water and ethylic alcohol 95%, the teeth were dried at room temperature and exposed to UV radiations in a cross-linker for 10 minutes on each side, in order to remove as much surface contamination as possible. Clean and irradiated teeth were submitted to a minimally destructive protocol to extract ancient DNA^107,108^ from the cellular cementum of tooth roots. The petrous bones were drilled to obtain 0.02 mg of bone powder, to be treated using the same protocol for the teeth.

The resulting DNA extracts were used to build paired-end, partially-UDG-treated, double-indexed single-stranded genomic DNA (ssDNA) sequencing libraries.^109^ All DNA extraction and library preparation steps were carried out in UCSC-PGL cleanroom facilities, following stringent contamination-prevention measures.^110^ To balance authenticity with data preservation, all libraries underwent “half-UDG” treatment, which partially removes deaminated bases from ancient DNA fragments.^111^ The prepared libraries were sequenced on Illumina NovaSeq 6000 platforms, generating 2×150 bp paired-end reads.

### Sequencing Reads Alignment

We used AdapterRemoval^112^ to trim adaptors from the sequence reads and collapse forward and reverse sequencing reads into single sequences, using the following parameters: --minlength 30 --minquality 25 --trimns --trimqualities. Alignment to the human reference genome (hg19) was performed using BWA^113^ (version 0.7.17) with the mem -k19 -r 2.5 parameters, following the suggestions of Xu et al.^114^, and SAMtools.^115^ Reads with a mapping quality below 30 were filtered out, and duplicate reads with identical orientation, start, and end positions were removed, retaining only the read with the highest sequence quality using DeDup.^116^

### Ancient DNA Data Authentication and Pseudo-haploid Variant Calling

We first calculated the C → T substitution rates for each individual using mapDamage^117^ to verify the authenticity of the ancient DNA. To control for post-mortem damage caused by spontaneous deamination of cytosines, we trimmed the three terminal bases on both sides of the reads. Next, we used Schmutzi^118^ to assess mitochondrial contamination by aligning the reads to the mitochondrial genomes of present-day individuals from diverse global populations. Any DNA libraries with insufficient data to estimate contamination were excluded from the downstream population genetic analysis (Table S2). Male individuals were tested for X-chromosome contamination using ANGSD.^119^

We used pileupCaller (https://github.com/stschiff/sequenceTools) to call variants at the 1.2 million genome-wide SNPs included in the 1240K panel.^44,45,120^ Any individuals sharing fewer than 10,000 SNPs with AADR (version 54.1),^44,45,120^ were excluded from the population genetic analyses (Table S2). We selected pseudohaploid data, generated by randomly sampling one allele per variant site, to reduce biases associated with low-coverage sequencing.

### Determination of Sex and Uniparental Haplogroups

We determined the biological sex of each individual by comparing the number of reads mapped to the Y-chromosome with those mapped to the autosomes or X-chromosome, using only high-quality reads with a mapping quality of 30 or above.^121^ The assignments were further confirmed through a derived pipeline that is also able to detect the chromosomal aneuploidy in ancient genomes.^122^ Specifically, genetic sex was inferred using two complementary approaches: the ratio of reads mapping to the X and Y chromosomes (R_y^121^) and chromosome coverage patterns estimated with the Karyo method^122^. Individuals with R_y < 0.016 were assigned female and those with R_y > 0.075 male, while karyotype was independently assessed from normalized X and Y chromosome coverage relative to autosomes. Assignments were considered reliable when based on ≥1,000 reads mapping to the sex chromosomes.

For the mitochondrial genomes, we constructed consensus sequences from reads that were at least 30 base pairs (bp) long and had a mapping quality of 30 or higher. We then assigned mitochondrial haplogroups to each individual using Haplogrep2,^123^ based on Phylotree Build 17.^124^ For male individuals, Y-haplogroups were identified using Yleaf (v3.1),^125^ focusing on sequences with both mapping and base qualities of 30 or higher.

Details about the molecular sex, mtDNA and Y-haplogroups for each individual can be found in Document S1, Table S2, and Figure S6.

### Biological Kinship Inference

We used READ^126^ to estimate the degree of biological relatedness across the sampled individuals from POG sharing at least 10,000 variants with AADR.^44^ This tool first computes the proportion of non-matching alleles (P0) between each pair of individuals, then normalizes it using the average P0 of unrelated individuals within a given dataset. The resulting normalized P0 values are used to categorize kinship relationships, such as whether two individuals are unrelated, first-degree relatives, second-degree relatives, or identical twins. The kinship results are shown in Table S3 and discussed in depth in Document S1. See also Figure S7.

### Dataset Compilation

We gathered previously published ancient (n = 642) and present-day (n = 974) human datasets for the 1240k SNP panel^1,15,32,120,127–140^ (Table S4) and combined them with the 18 unrelated newly sampled ancient individuals from POG. For the present-day data, we incorporated genome-wide data from Western Eurasia (Italy, Spain, UK, France, Germany) and Africa (Yoruba). Similarly, we included ancient individuals focusing on North Sea facing areas.^1,140^

### Principal Components Analysis

For the Principal Component Analysis, we combined both ancient and modern samples into a dataset using the 1240k SNP panel, and then projected the ancient samples into the principal component space based on modern Eurasian populations. We used the smartpca program from the EIGENSOFT package^141,142^ to carry out the analysis, with default settings except for the “lsqproject: YES” option. We initially included present-day humans from across Eurasia. Then, we projected both our newly sequenced and previously published ancient samples into this space.

### Population Structure Analysis

We used ADMIXTURE^143,144^ to explore the population structure of POG considering the newly sequenced unrelated individuals, together with 974 present-day and 642 ancient samples. For this analysis, we removed variants in moderate to high linkage disequilibrium (r² > 0.4), SNPs with more than 60% missing data, and those with minor allele frequencies below 5%. This was done using PLINK^145^ with the parameters “--geno 0.6 --maf 0.05 --indep-pairwise 200 50 0.4” leaving us with 75,048 SNPs. To ensure balanced representation of ancient populations, we analyzed a restricted dataset limited to a maximum of 15 individuals per group, resulting in a total of 241 samples. Among these, three groups were included as potential outgroups to enhance resolution by polarizing allele frequencies: *France_A* (Iron Age individuals from the Marne region in northeastern France), *Serbia_A* (Roman-period individuals from Serbia), and *Italy_A* (Romans from central Italy). We ran the ADMIXTURE unsupervised analysis for K values ranging from 2 to 10. Since K = 4 had the lowest CV error (Figure S4), we used this value for our final ADMIXTURE results, which are shown in Figure 2c. A supervised approach followed the unsupervised runs to yield more precise ancestry estimates considering Saxon as a single source, comprising *UK_Sx*, *Denmark_Sx*, *German_Sx*, and *Netherlands_Sx*, while Viking groups were distinguished based on their geographical designations in *Sweden_Vk*, *Denmark_Vk*, and *Norway_Vk*. Moreover, a French source was considered, merging *France_LA*, *France_GalloRom*, and *France_A*.^144^

### Qutgroup-f3 and D-statistics

To assess the genetic similarity between pairs of populations, we calculated the outgroup-f3 statistic using the qp3Pop software (v412) from the AdmixTools package.^146–148^ The statistics were calculated in the form of f3(Yoruba; X, Y), with the present-day Yoruba population from Central/Western Africa serving as the outgroup, after processing their data using “--geno 0.4 --maf 0.05 --indep-pairwise 200 50 0.4” in PLINK.^145^ We also used D-statistics to explore the genetic relationships among four populations, employing the qpDstat software (v712) from the AdmixTools package.^146–148^ This was done for both the newly sampled POG individuals and several published ancient and present-day populations, with the D-statistics calculated in the form of D(P1, P2; P3, Outgroup).

### Admixture Modeling

We used qpWave from Admixtools library in R and qpAdm (version 634) from the AdmixTools package^146–148^ via AdmixR^115^ to model the ancestry proportions of the POG population. First, we extracted f2 blocks and then we tested for putative sources the Northern Europe ancient population other than POG. The outgroups group comprised: ‘YRI’, ‘Italy’, ‘Spain’, ‘ALG’, ‘Italy_A’, and ‘Serbia_A’, for which recent admixture with POG can be excluded. We tested different mixture models involving one and two source populations. To minimize the impact of missing data in some populations, we used the “allsnps: YES” parameter.^146^ Once a significant model was built in qpWave, we used the populations in the models for determining their relative contribution to POG. We used *models2components()* function in AdmixR library to rotate the factors and identified good-fit models related proportions according to p-value > 0.05, and ancestry proportions > 0.

## ACKNOWLEDGEMENTS

Research reported in this publication was supported by the National Institute of General Medical Sciences of the National Institutes of Health under Award Number R35GM142939 for C.E.G.A. The content is solely the responsibility of the authors and does not necessarily represent the official views of the National Institutes of Health. Additional support was provided by start-up funds from the College of Science and Mathematics at California State University, Northridge, to C.E.G.A. F.D.A. was partially supported by MNESYS: a multiscale integrated approach to the study of the nervous system in health and disease (PNRR). The dietary isotopes and collagen extraction were funded by a Cambridge Trust and Newnham College Scholarship for S.L. (no. 10386281). Radiocarbon dating was funded as a National Environmental Isotope Facility grant-in-kind (NEIF 2438.1021) for L.B., T.C.R., S.L, and R.P. The following grants and fellowships funded and supported the radiocarbon modelling for this research all awarded to S.L.: a Leverhulme Trust Early Career Fellowship (ECF-2021-467), a University of Edinburgh School of History, Classics and Archaeology Research and Travel grant and a University of Edinburgh Centre for Data, Culture and Society Training Bursary. C.E.G.A. thanks Tábita Hünemeier (Institut de Biologia Evolutiva, Consejo Superior de Investigaciones Científicas, Spain) for helpful discussions. S.L. thanks Catherine Kneale (McDonald Institute, Cambridge) for her assistance with isotopic sample preparation and analysis via EA-IRMS. We also wish to thank the many students who volunteered their time over several summers to help clean and study the remains for the osteological analyses.

## AUTHOR CONTRIBUTIONS

Conceptualization: F.D.A., S.L., L.B., and C.E.G.A. Genomics experimental work: F.D.A., E.A.N., N.B., K.K., and L.F.-S. Genomics laboratory supervision: L.F.-S. Isotopes experimental work: S.L. Curation of the human remains and osteological analyses: T.C.R. and L.B. Archeological and historical contextualization: S.L. and R.P. Data curation and formal analysis: F.D.A. (genomics) and S.L. (isotopes). Visualization: F.D.A. and T.R.P. Funding acquisition: S.L., L.B., and C.E.G.A. Project administration and supervision: C.E.G.A. Writing-original draft: F.D.A. and C.E.G.A. All authors reviewed the final version of the manuscript.

## DATA AVAILABILITY

The accession number for the sequencing data reported in this paper is [PLACE HOLDER], deposited in the European Nucleotide Archive (ENA). FASTQ files containing endogenous human are provided for each individual. Metadata, sequencing statistics, contamination estimates, and uniparental haplogroup assignments for each individual are provided in Table S2. Additional information required to reanalyze the data reported in this paper is available from the corresponding authors (F.D.A. and C.E.G.A. for the genomic data; L.B. and C.E.G.A. for anthropological and archeological data) upon request.

## SUPPLEMENTAL INFORMATION

**Document S1**. Figures S1-S7 and supplemental references

Data S1. OxCal script used for mixed marine-terrestrial calibration and modeling of radiocarbon dates from Priory Orchard of Godalming (POG).

**Table S1. Stable isotope and radiocarbon data for individuals from Priory Orchard of Godalming (POG).** This spreadsheet contains three tabs presenting isotope and radiocarbon data. Tab 1 (“dbExport_C14_dates”) provides raw stable isotope measurements (δ¹³C and δ¹□N) for human remains together with faunal end-member values. Tab 2 (“POG_dates_uncertainty_forimport”) reports raw, uncalibrated radiocarbon determinations (¹□C BP) with associated laboratory errors/uncertainties. Tab 3 (“%marinecalcs”) presents calibrated radiocarbon dates modeled in OxCal v4.4 using mixed marine and terrestrial calibration curves, including calibrated age ranges and summary statistics used in the chronological analyses described in the manuscript.

**Table S2. Sequencing statistics, authentication metrics, radiocarbon dating, and metadata for Priory Orchard of Godalming (POG) genomic libraries.** Sequencing and mapping statistics for all Priory Orchard of Godalming (POG) libraries screened for ancient DNA (aDNA). “Skeletal ID” refers to the archaeological sample identifier and “Study ID” to the laboratory code. Skeletal ID should be used for references to the data presented in this study. “Dating LB” and “Dating UB” indicate the lower and upper bounds of the calibrated radiocarbon date (mixed calibration curve, 95.4% probability), while “Dating” represents the mean calibrated date. “Osteo Sex” corresponds to osteological sex estimation. Sequencing metrics include “Reads count” (total sequencing reads), “Mapped” (mapped reads after duplicate removal), “Endo (%)” (percentage of endogenous DNA), “Mean cov” (mean genome-wide coverage), and “1240k cov” (average coverage on the 1240K SNP panel). Contamination estimates are reported as “X-chrom cont” (X-chromosome contamination estimate) and “Mito cont” (final endogenous mitochondrial contamination estimate from Schmutzi). Sex determination using the R_y method includes “Nseqs” (number of reads used for sex determination), “NchrX+NchrY” (reads mapping to chromosomes X and Y), “NchrY” (reads mapping to chromosome Y), “R_y” (ratio of Y-chromosome reads), “SE” (standard error), and “95% CI” (95% confidence interval), followed by the resulting assignment. Sex determination using the karyo_RxRy method includes “Na” (number of reads mapping to autosomes, excluding chromosomes 13, 18, and 21), “Rx” (proportion of reads mapping to chromosome X over Na), “RxSE” (standard error of Rx), “Ry” (proportion of reads mapping to chromosome Y over Na), “RySE” (standard error of Ry), and the resulting assignment. “Sex” indicates the final karyotype attribution based on both R_y and karyo_RxRy methods. Uniparental markers include “mt_coverage” (mitochondrial DNA coverage), “Ychr Hg” (Y-chromosome haplogroup), and “mtHg” (mitochondrial haplogroup assignment). “Quality” indicates haplogroup assignment quality as assessed by Haplogrep3, and “Range” indicates the mtDNA regions used for haplogroup determination. Post-mortem damage patterns are reported as “1 pos 5pC>T”, “2 pos 5pC>T”, and “3 pos 5pC>T”, representing the frequency of C→T substitutions at positions 1, 2, and 3 from the 5′ end of sequencing reads, respectively.

**Table S3. Pairwise kinship relationships among POG individuals.** Pairwise relatedness estimates among POG individuals inferred using READ (see STAR Methods). The table reports the inferred degree of relatedness (e.g., First Degree, Second Degree, Identical Twins, Unrelated). “Z_lower” and “Z_upper” represent the Z-scores relative to the lower and upper classification thresholds, respectively. The column “SNP” returns the total number of SNP windows used in each estimation.

**Table S4. Population panels used in ancestry analyses.** Tab S4A lists the population panel derived from the AADR v54.1 dataset. Tab S4B lists the subset of samples (n = 241) constituting the restricted panel used for ADMIXTURE analyses. Associated metadata were obtained from AADR v54.1.

**Table S5. Results of qpWave analysis**. “Pop1” indicates the test population and “Pop2” the comparison population. “Pop3” and “Pop4” represent outgroups 1 and 2, respectively. “Est” denotes the f4-statistic estimate, “SE” the standard error, “Z” the Z-score, and “P” the associated P-value.

**Table S6. Results of qpAdm ancestry modeling.** Plausible models (i.e., models with both ancestry proportions > 0 and P > 0.05) are highlighted in bold. “Model” indicates the model identifier, “Target” the target population, “Source1” and “Source2” the putative source populations, and “Outgroups” the set of outgroup populations used in the model. “Noutgroups” indicates the number of outgroups included. “P-value” represents the statistical fit of the model. “Proportion_pop1” and “Proportion_pop2” denote the estimated ancestry proportions attributable to Source1 and Source2, respectively, with “stderr1” and “stderr2” indicating their corresponding standard errors.

