## Supplemental Document S1 for "The Genomic Legacy of the Norman Conquest in Rural England"

**SUPPLEMENTARY INFORMATION**

**Historical and Archaeological Context**

The North Sea and English Channel acted as major conduits of human mobility, trade, and cultural exchange throughout the first millennium CE, linking the British Isles with Scandinavia, the Low Countries, and France.^1,2^ Archaeological evidence indicates sustained interaction across these regions, including shared material culture such as brooches, combs, pottery styles, and metalwork, as well as similarities in burial practices, architectural features, and coinage circulation^3–6^. These exchanges reflect long-term, structured networks of contact, rather than sporadic or incidental movement. This is in part due to Scandinavian settlements in Britain, Ireland, and the rest of Europe during the Viking Age, which began in the mid 8^th^ century. The first documented raid on England was in Portland, Dorset, in 789 CE.^7^ By 865 CE, the ‘Great Army’ had reached Britain and, in its wake, the Danelaw was established in eastern and northern England, with upwards of 35,000 settlers estimated by the end of the 10^th^ century.^8,9^ Indeed, archaeological evidence from Viking-period settlements in eastern and northern England further supports this pattern, showing a mix of Scandinavian-style dwellings and material culture embedded within predominantly Anglo-Saxon landscapes. These observations demonstrate that “Viking” identity in England was largely cultural, reflecting networks of migration, social integration, and economic participation, rather than from large-scale demographic replacement.^10^

Viking-age movements exemplify the complexity of North Sea interactions. Although historical sources emphasize raiding, pillaging, and military conquest, genomic and isotopic evidence indicates that Viking-associated groups also engaged extensively in settlement, social integration, and intermarriage with local populations.^10–12^ Isotopic analyses of early medieval skeletal remains document both local continuity and non-local origins, revealing mobility patterns shaped by voluntary migration, interregional marriage, trade, and, in some cases, displacement associated with conflict. For example, strontium isotope data from eastern English cemeteries indicate that a subset of individuals originated in continental Europe, underscoring the heterogeneous composition of rural populations even outside major urban centers^13,14^. This complexity is further illustrated by genomic analyses of mass burials such as those in St. John’s College (Oxfordshire, ca. 1000 CE) and Ridgeway Hill (Dorset, 970 - 1025 CE), which reveal heterogeneous ancestry profiles, with individuals carrying both Scandinavian-like and local Anglo-Saxon components.^15,16^ These findings are consistent with isotopic evidence of variable childhood origins, suggesting that some individuals were born locally while others migrated from Scandinavia^17^.

At broader regional scales, additional differences become apparent: Danish-influenced groups predominated in the Danelaw regions of eastern and northern England, whereas Norwegian-like groups were concentrated in the Irish Sea zone and northern Britain, contributing to geographically stratified genetic landscapes.^10,18,19^ Genomic analyses further show that Danish-like ancestry within those groups partially overlapped with that of ‘Anglo-Saxon’ individuals, which is not surprising given their proposed origins.^10,20,21^ The influence and overlap between England and Denmark in particular was such that in the 11^th^ century there were four Danish kings of England (Sweyn Forkbeard, Cnut, Harold Harefoot, and Harthacnut) and under Cnut both countries were governed as part of the North Sea Empire.^22^

It is through these Danish connections that the Norman claim on England originated.^23–25^ The Normans, who descended from Norse settlers in Northern France in the 9^th^ century (with *Normandy* derived from the Old French for ‘North man’), blended Norse, Frankish, Breton, and Gallo-Roman influences into a distinct identity which was consolidated under the new Duchy of Normandy in 911 CE.^22,26^ The Norman presence at Edward the Confessor’s court thus exerted influence long before the Conquest of 1066; However, it was only after the Conquest that most social, political, and cultural changes took place.^26–29^ Following William I’s victory in October 1066, the Normans swiftly consolidated power, implemented widespread administrative reforms, and instigated the Domesday Survey.^30^ The Norman Conquest reshaped England’s political, administrative, and social structures, including the introduction of feudal governance, redistribution of land to Norman lords, establishment of new ecclesiastical hierarchies, and the construction of stone castles and manorial estates. Archaeologically, the Norman presence is more clearly visible in urban centers and aristocratic contexts, where imported luxury goods, Romanesque architecture, and altered material culture appear abruptly.^30–32^ In contrast, rural communities, particularly in the south, such as those represented at Priory Orchard of Godalming (POG), likely experienced more gradual demographic and cultural shifts, while retaining continuity in settlement patterns, subsistence practices, and burial customs.^31,32^ For example, the presence of standardized supine inhumations without significant changes in grave goods at POG suggests cultural continuity alongside subtle adoption of Norman-influenced practices. Isotopic and genetic data from such rural cemeteries are therefore essential for identifying localized demographic impacts, which may differ substantially from those inferred from urban or elite contexts.

Southern England from the 9^th^ through the 12^th^ centuries was shaped by successive and overlapping waves of migration and cultural influence across the North Sea and English Channel. Interactions among Anglo-Saxon, Viking, and Norman populations generated a complex mosaic of ancestry, cultural identity, and local integration in England as a whole. Genomic analysis of rural sites like POG enables direct assessment of these dynamics, providing insight into how large-scale political events -- such as the Norman Conquest -- translated into population-level genetic outcomes, while also illuminating patterns of continuity, admixture, and social incorporation in rural medieval communities.

**Sex Chromosome Karyotype Inference and Uniparental Markers**

Sex Chromosome Karyotype Inference

To explore the demographic features and descent practices of populations from Medieval Surrey, we examined the sex assignments and uniparental haplogroups of the 78 POG individuals. Among them, 21 females and 17 males were identified, while 40 osteological remains returned insufficient read counts for the biological sex assignment analysis (Table S2).^33^

Four male individuals (POG14-1033, POG14-1079, POG15-3148, and POG15-3258) were identified as potential carriers of a chromosomal aneuploidy: an additional copy of chromosome Y (47,XYY karyotype), which is currently observed in about 1 in 1000 male births.^34^ We note that a similar observation was made for an Early Medieval Period (8^th^ century) archaeological site from the Lincoln Eastern Bypass, Lincoln.^33^ However, the low depth of coverage for POG14-1079 (0.011x), POG15-3258 (0.037x), and POG15-3148 (0.061x) makes these results unreliable for these three individuals. Conversely, POG14-1033 (0.867x) is in line with the coverage obtained from the aforementioned individuals from Lincoln Eastern Bypass, prompting broader consideration of the prevalence of sexual aneuploidies in medieval England.^33^

Mitochondrial DNA Lineages

The mitochondrial haplogroup composition of this medieval British population reveals a **remarkably diverse and stratified maternal ancestry**, reflecting long-term population continuity in the British Isles as well as the cumulative effects of repeated demographic events from the Mesolithic through the Middle Ages. The predominance of haplogroup H and its multiple subclades (Figure S6) – encompassing a wide array of subclades including **H1, H1a, H1c1, H1c13, H1be, H1g1, H1ai1, H3+152, H45b, H53, H5b, H6a1a, H6a1b4, and H2/H2a/H2a2** – reflects lineages widely distributed across Europe since the post-glacial period and Neolithic expansion,^35^ indicating the persistence of established European maternal lineages in Britain through time. The frequencies of these lineages are consistent with other medieval and earlier European populations, suggesting continuity of broad European maternal lineages in Britain across time.^36,37^

Haplogroups K, T, and J, commonly associated with Neolithic farming populations,^38–40^ indicate the lasting genetic contribution of early agriculturalists. The presence of multiple subclades, such as **K1a, K1a4a1a+195, K1a4a1e*, K1b2b, K1e, K2a, T1a1, T2b5, T2b5a1, T2b9, T2e1a1b, and J2a1a1a2**, indicates that the genetic legacy of early agriculturalists remained a **stable and enduring component of the maternal gene pool** well into the medieval period. Specifically, haplogroup J has been linked to the expansion of agriculture from West Asia into Europe and later into Scandinavia,^38^ where multiple J sub-haplogroups are present by the Viking Age; its occurrence in medieval Britain may therefore reflect both early Neolithic ancestry and later population interactions.

The detection of **U4, U5, and U8** haplogroups – including **U4a2, U5a1a2b, U5a2a1d, U5b2b4a, U5b2b3a1a, and U8 –** underscores the persistence of **Mesolithic hunter-gatherer maternal lineages** within the POG population. These haplogroups represent some of the oldest mitochondrial lineages in Europe and are indicative of deep ancestry.^41,42^ Their continued presence in medieval Britain suggests that, despite major demographic transitions such as the Neolithic and later migrations, **maternal lineages associated with Britain’s earliest inhabitants were not eliminated but instead integrated into later populations** through admixture and continuity. This finding aligns with broader ancient DNA evidence showing substantial survival of hunter-gatherer maternal ancestry in northern Europe.^37^

The presence of less common mitochondrial lineages, such as V, W3a1, X2c1a, and I4a, further underscores the maternal heterogeneity observed at POG. These haplogroups occur at low frequencies across a wide range of European populations and are not readily attributable to any single migratory episode or cultural group.^43,44^ Their occurrence in the POG assemblage likely reflects the cumulative effects of long-term population mobility, gene flow, and social integration operating over multiple periods, rather than discrete historical events. Together, these rare lineages highlight the complex and layered maternal ancestry of this rural medieval population, consistent with sustained connectivity across Europe over millennial timescales.

Taken together, the mitochondrial haplogroup diversity observed in POG reflects a layered maternal genetic structure shaped by deep prehistoric ancestry, Neolithic agricultural expansions, and later historical population interactions. The predominance of haplogroup H indicates continuity of broad European maternal lineages, while the substantial representation of K, T, and J lineages points to lasting Neolithic influences, and the persistence of U lineages highlights the survival of Mesolithic-associated ancestry. The presence of rarer haplogroups further underscores the heterogeneity of the maternal gene pool, consistent with long-term population connectivity rather than discrete migratory events. Collectively, these patterns indicate that the maternal genetic makeup of POG reflects the cumulative outcome of long-term continuity and repeated admixture over multiple periods, rather than the result of a single demographic episode.

Y-chromosome Haplogroup Diversity

The Y-chromosome haplogroup profile observed in POG (Table S2) reflects a layered paternal ancestry, resulting from the complex demographic processes that shaped early and medieval England. The predominance of R-M269 and its subclades (R-L23*, R-YP356*, R-Y30815*, R-Z30*, R-Y179346*) highlights a major contribution from populations associated with the Anglo-Saxon migrations.^45^ Historical and genetic evidence suggests that Saxon settlers, moving from continental northern Europe between the 5^th^ and 7^th^ centuries CE, established a substantial male presence in England, and the widespread presence of R-M269 today indicates that this lineage became numerically dominant, forming a foundational component of the paternal gene pool.^46,47^ The presence of R1a* and R-Y11145* reflects the influence of Viking populations, particularly from Scandinavia, who settled in the Danelaw and surrounding regions during the 9^th^ to 11^th^ centuries CE.^48^

Haplogroups I-Y7219* and I1* represent Northern European and Germanic lineages, encompassing both pre-Saxon inhabitants and migrants such as the Franks, who had intermittent contact with England during the early medieval period.^49^ These lineages occur at moderate frequencies and likely reflect additional paternal diversity alongside the dominant R-M269 background, consistent with a mixture of long-standing local lineages and broader continental European gene flow.

The detection of J-Y294088*, a branch of haplogroup J,^50^ could suggest a minor paternal contribution from the Mediterranean or Near East, potentially introduced through long-distance trade networks, migration of elites, or other limited gene flow during the Middle Ages. However, the low number of reads mapping on the Y chromosome for this individual makes this assignment only tentative.

Overall, the Y-chromosome composition suggests a hierarchy of male ancestry in POG, dominated by putatively Saxon-derived R-M269 lineages, with secondary contributions from Viking and continental Germanic populations, and only minor inputs from southern European or Near Eastern sources. This pattern might mirror both historical records of migration and settlement and the genetic landscape observed even in present-day populations of the British Isles, illustrating how successive waves of migration and regional interactions shaped the paternal gene pool over centuries.

**Biological Kinship Inference**

The biological relatedness (kinship) analysis indicated highlighted several related pairs, including 26 first-degree relationships – such as parent-child or full siblings – and 43 second-degree kinships, involving grandparents, grandchildren, or aunt/uncle-niece/nephew relationships (Table S3). We note that these relatedness estimates may also capture more complex pedigree configurations or elevated background relatedness within the population, and therefore do not uniquely specify a single genealogical relationship.

Additionally, two pairs were classified as identical twins or representing the same individual, despite originating from different excavation areas. Specifically, these pairs are POG14-1079 and POG15-3107, and POG14-1041 and POG15-3102 (Figure S7).

POG14-1079 and POG15-3107 are both dated to approximately 1015 CE (Table S1). Sequencing coverage differs markedly between the two individuals (0.011× and 0.465×, respectively), and their sex assignments are discordant both osteologically and genetically (Table S2), as are their mitochondrial haplogroup calls (T2b5 and T1a1, respectively). We note, however, that the mtDNA coverage for POG14-1079 contains substantial gaps, rendering the haplogroup assignment for this individual uncertain. In addition, POG15-3107 was inferred to be first-degree related to 12 individuals with substantially higher genomic coverage (1240K SNP coverage ≥ 10,000 SNPs) buried in different areas of the POG site (Table S3). In contrast, relatedness between POG14-1079 and these same individuals could not be reliably assessed due to the limited number of overlapping SNPs. In sum, while some of these observations could be consistent with POG14-1079 and POG15-3107 representing twins, the discordant sex assignments and mitochondrial haplogroups, together with their differing patterns of relatedness to other individuals, argue against a definitive identification. As such, this interpretation remains unlikely, or at best tentative.

The second putative twin pair (POG14-1041 and POG15-3102) consists of individuals with discrepant dating estimates (Table S1), centered at approximately 1015 CE and 1053 CE, respectively, but with largely superimposing confidence intervals. Osteologically, their estimated age at death is 25-35 and 17-25, respectively. Sequencing coverage differs substantially between the two individuals (0.784× and 0.024×, respectively), and such pronounced differences in data quality can compromise kinship inference. In addition, sex assignment for POG15-3102 remains inconclusive due to limited sequencing coverage, including of the sex chromosomes. Although both samples exhibit sufficient mitochondrial coverage for haplogroup assignment, sequencing gaps may account for the observed differences in their macro-haplogroups (H6a1a and H53, respectively). Taken together, these factors suggest that the inferred close kinship between POG14-1041 and POG15-3102 is uncertain and may reflect limitations of the data rather than a true biological relationship.

**Supplementary Figures**

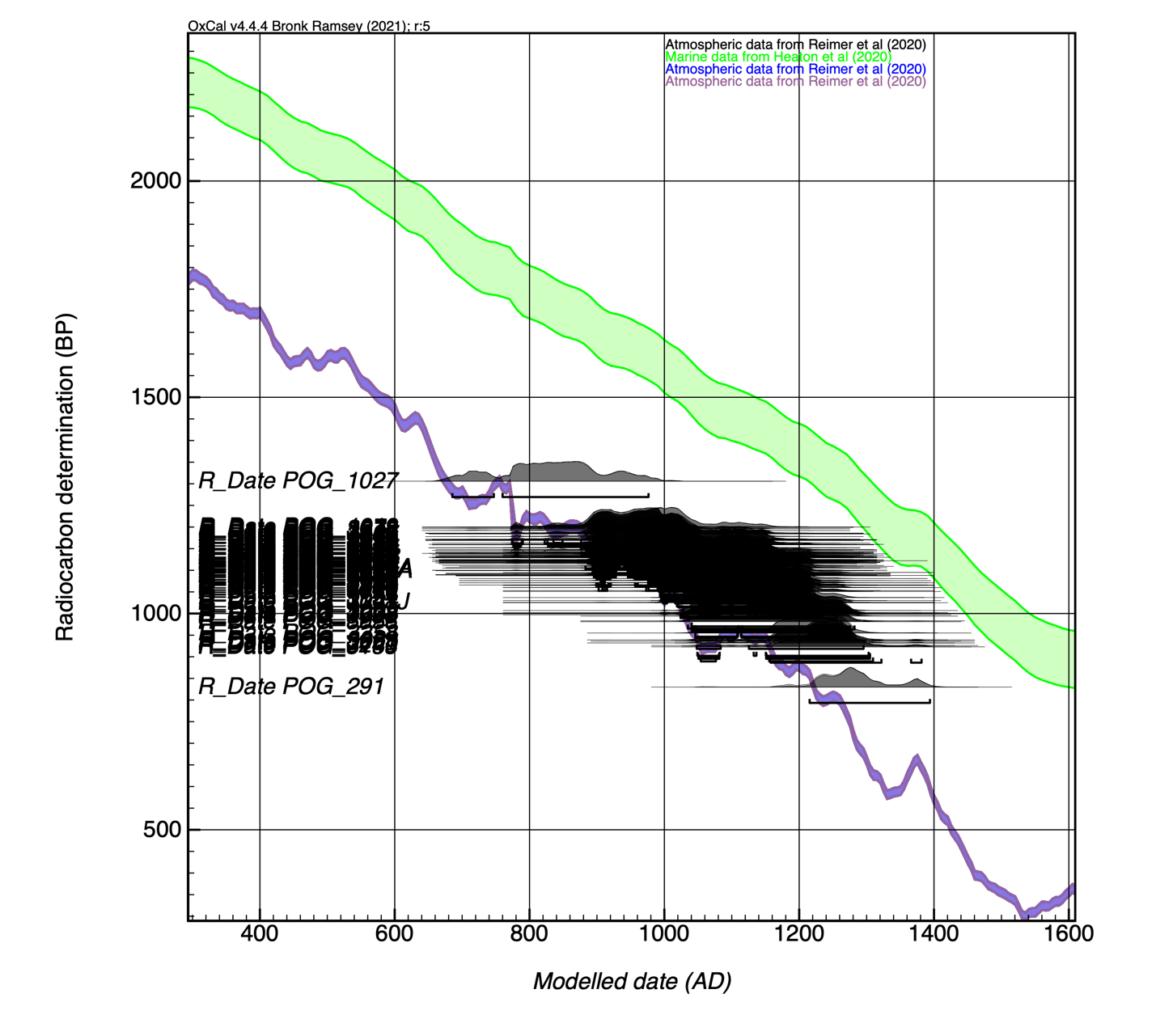

**Figure S1: Mixed-curve calibrated radiocarbon dates for human remains from Priory Orchard of Godalming (POG), related to STAR Methods.** Calibrated radiocarbon probability distributions for POG individuals modeled in OxCal v4.4 using mixed marine and terrestrial calibration curves. Each distribution reflects the calibrated age probability range for an individual sample (see also Table S1).

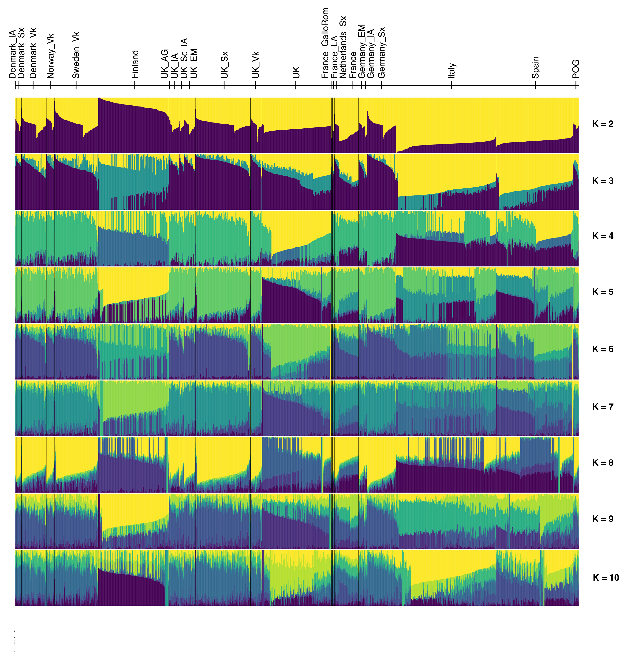

**Figure S2: Model-based clustering results across multiple values of K, related to Figure 2.** Unsupervised ADMIXTURE clustering for the merged dataset (n = 1,634 individuals; Table S4A) across values of K = 2–10. Each vertical bar represents an individual and colors correspond to inferred ancestry components. Individuals are grouped by population and ordered consistently across runs, illustrating the structure of ancient and present-day populations included in the analysis.

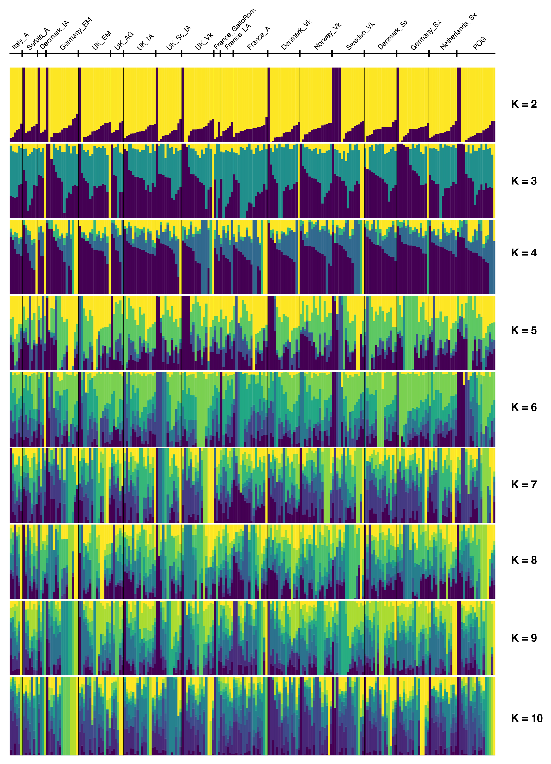

**Figure S3: Model-based clustering results across multiple values of K, related to Figure 2.** Unsupervised ADMIXTURE clustering for the restricted dataset (n = 1,634 individuals; Table S4B) across values of K = 2–10. Each vertical bar represents an individual and colors correspond to inferred ancestry components. Individuals are grouped by population and ordered consistently across runs, illustrating the structure of ancient and present-day populations included in the analysis.

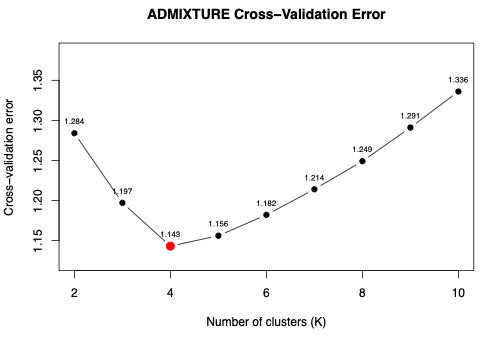

**Figure S4: Cross-validation error across ADMIXTURE models, related to Figure 2.** Scree plot showing ADMIXTURE cross-validation (CV) error across different numbers of ancestral clusters (K). CV error values are shown for K = 2–10, with lower values indicating better model fit.

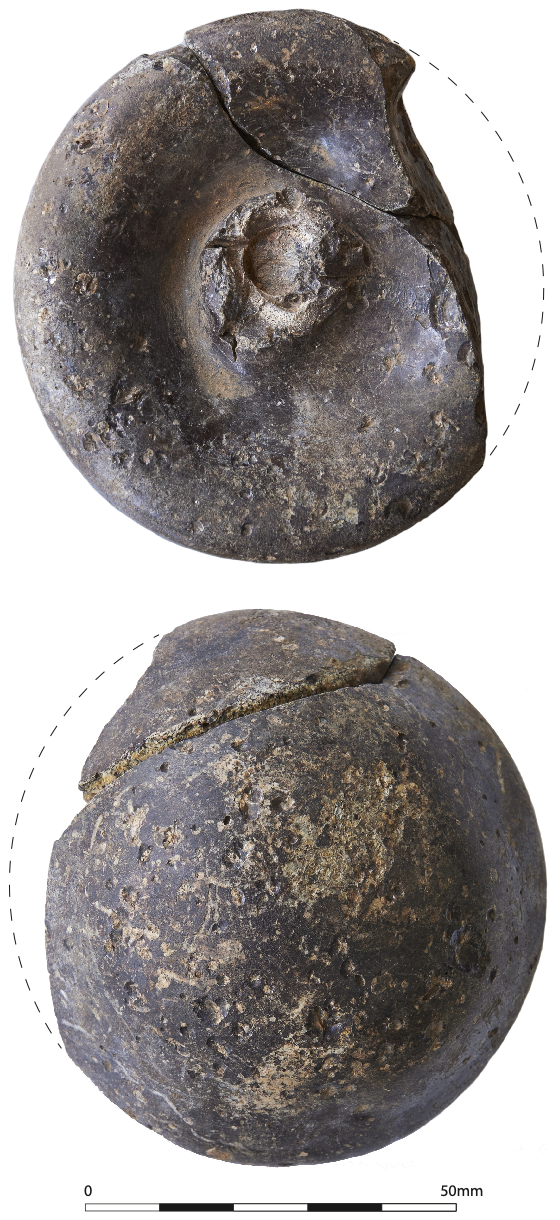

**Figure S5: Glass linen smoother carefully positioned over the right midriff of burial 3107 in the cemetery at Priory Orchard, Godalming, related to Figure 3.** This object was one of only three items intentionally placed with burials in the cemetery and may have marked the individual as someone of particular status; similar smoothers are known from Viking-age contexts in Britain and Scandinavia, where they are often associated with female burials and interpreted as indicators of high social standing. In that perspective, it should be borne in mind that early medieval glass smoothers have also been reconsidered as tools used for gilding Christian murals rather than primarily for smoothing linen fabric. These circular glass objects, discovered across various parts of Europe, contain high concentrations of metals such as gold, silver, copper, zinc, or mercury, suggesting they may have also been employed to apply metal leaf onto church walls, potentially serving as a symbolic act associated with the dissemination of Christianity. Image copyright: Surrey County Archaeological Unit (part of Surrey County Council).

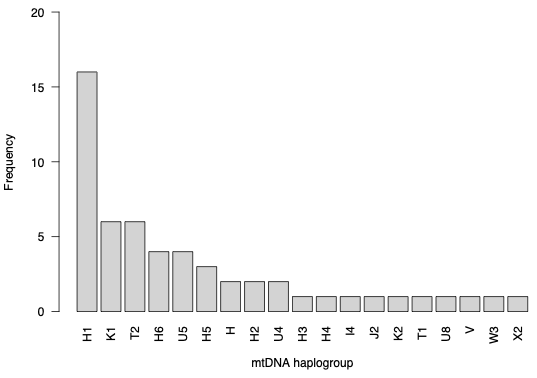

**Figure S6: Frequencies of mitochondrial DNA (mtDNA) lineages.** Frequencies (counts) of mtDNA haplogroups observed in the POG dataset. Minor clades are collapsed into major haplogroup designations (first two digits).

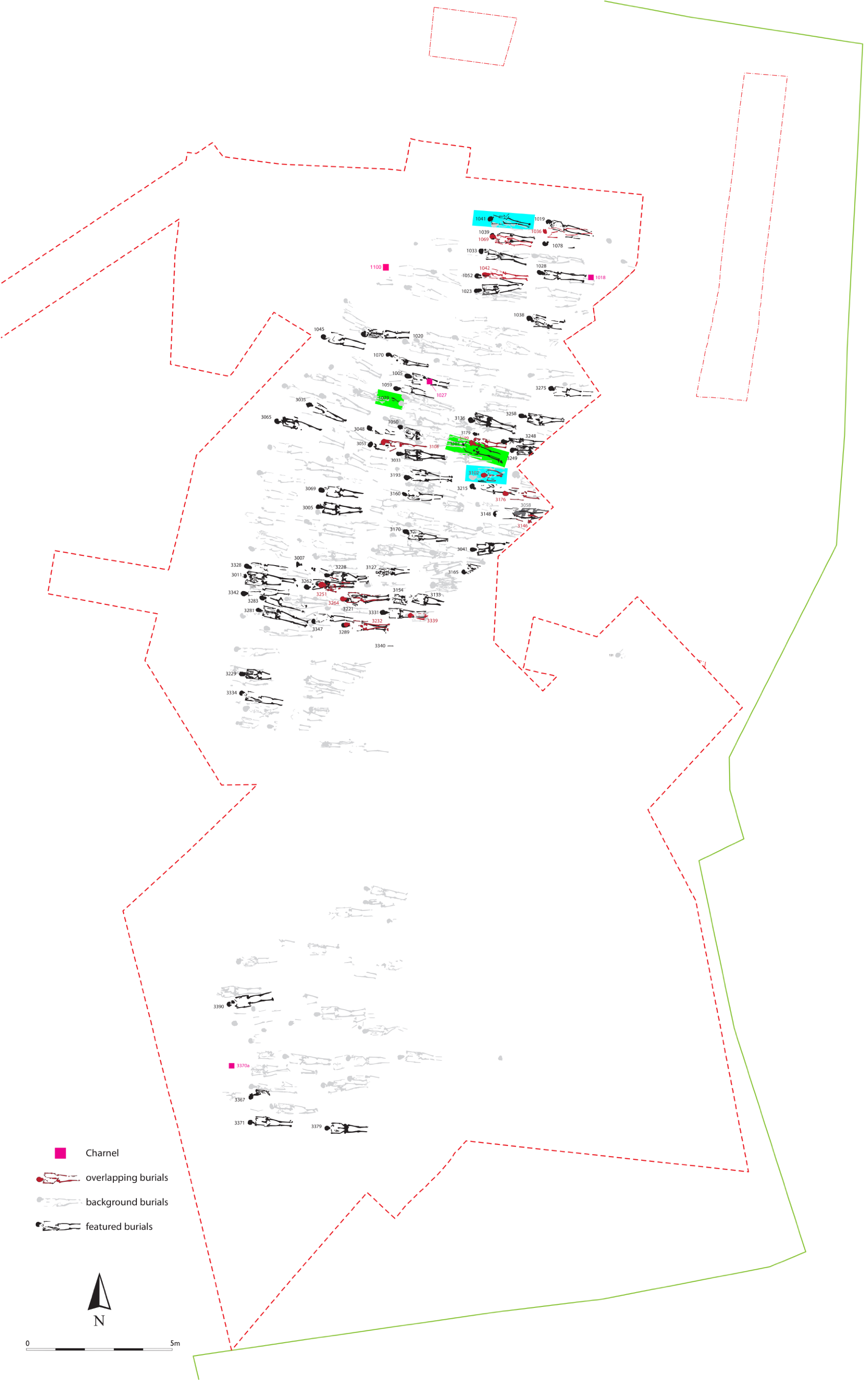

**Figure S7: Spatial distribution of burials, highlighting two pairs of individuals identified as identical twins.** Pairs of individuals classified as identical twins or representing the same individual are shown within the overall burial layout of the cemetery. The corresponding pairs are highlighted in color.
